# Antha-guided Automation of Darwin Assembly for the Construction of Bespoke Gene Libraries

**DOI:** 10.1101/2019.12.17.879486

**Authors:** P. Handal Marquez, M. Koch, D. Kestemont, S. Arangundy-Franklin, V. B. Pinheiro

## Abstract

Protein engineering through directed evolution facilitates the screening and characterization of protein libraries. Efficient and effective methods for multiple site-saturation mutagenesis, such as Darwin Assembly, can accelerate the sampling of relevant sequence space and the identification of variants with desired functionalities. Here, we present the automation of the Darwin Assembly method, using a Gilson PIPETMAX™ liquid handling platform under the control of the Antha software platform, which resulted in the accelerated construction of complex, multiplexed gene libraries with minimal hands-on time and error-free, while maintaining flexibility over experimental parameters through a graphical user interface rather than requiring user-driven library-dependent programming of the liquid handling platform. We also present an approach for barcoding libraries that overcomes amplicon length limitations in next generation sequencing and enables fast reconstruction of library reads.

## 1. Introduction

Large-scale screening and characterization of protein libraries is a powerful approach for both protein engineering and directed evolution as it enables (depending on the platform) sampling of up to 10^12^ [1] variants facilitating the identification of proteins with tailored functionalities [2,3,4,5]. Nevertheless, that represents only a miniscule fraction of the available variation in a small 100-residue protein (where 10^12^ variants equate to approximately 1 x 10^-116^% of the available sequences). Therefore, library design and assembly are powerful steps in which to maximize the relevant sampling space [6]. Numerous techniques for single- and multiple-point saturation mutagenesis exist with various degrees of efficacy (fraction of population that is mutated) and efficiency (fraction of mutant population where all targeted sites are mutated) [1,7,8]. We recently developed the Darwin Assembly (DA), which is a highly efficient and effective multiple-site saturation mutagenesis that can target up to 19 non-contiguous residues with less than 0.25% wild-type contamination. The assembly strategy was designed with automation in mind and here we demonstrate that its automation not only accelerates library construction but also enables the generation of highly complex libraries with minimal hands-on time and with few possibilities of human error.

The Darwin Assembly method presented here has been physically executed on a Gilson PIPETMAX^TM^ liquid-handling robot, controlled using a custom-built Element Suite in Antha; a third party software that facilitates high-level, biological-directed programming of liquid-handlers [9].

Antha is a software platform comprising experimental workflows encoded in a custom programming language (derived from Google Go), an experimental planner which defines resource requirements and program logic, device drivers for translating instructions to laboratory automation platforms (principally liquid handlers), a graphical interface, and an application for controlling the automated systems in use. Users of the system, using a graphical interface, construct workflows representing the flow of materials and data through their experiment. The users provide experimental parameters and data, which the software uses to generate instructions for a chosen set of devices. By applying methods from compiler design and optimization, Antha allows complex liquid handling workflows to be created directly from the outputs of the design tools, without the need for manually specifying the details of the large number of liquid transfers which that entails and that are part of the standard methods of programming such platforms.

The Antha DA automated workflow plans and physically executes all liquid-handling steps of the DA method (bar the magnetic bead purification steps), from the generation of complex mutagenic primer mixes to the preparation of recovery PCR reactions post-assembly – all 8 steps required in the assembly experimental pipeline (see Fig. 1). As examples of the platform, we report the assembly of 13 3-point mutant libraries and 38 2-point mutant libraries targeting a total of 39 residues in the *Thermococcus kodakariensis* DNA polymerase (KOD DNAP) gene for the screening of improved XNA synthesis. We also generate 16 single residue saturation mutagenesis libraries on the human proteasome subunit β5 (PSMB5) gene targeting a total of 48 residues. Finally, we also demonstrate a strategy to overcome read-length limitations of Next Generation Sequencing (NGS) by barcoding libraries and reconstructing entire length reads for the identification of all mutations.

**Figure 1.**
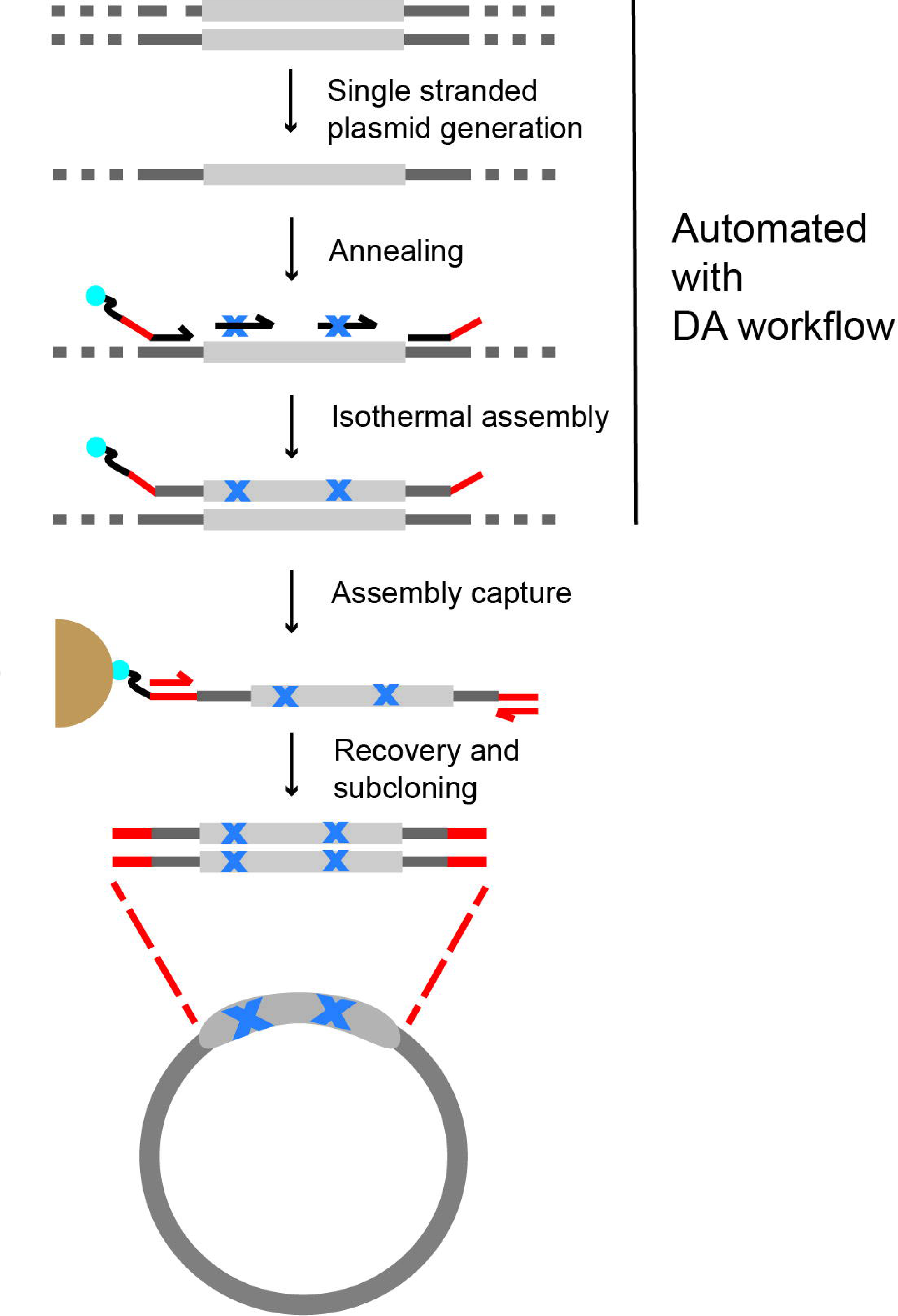
Darwin Assembly with biotinylated oligonucleotides. Schematic representation of Darwin Assembly with biotinylated oligonucleotides (adapted from Cozens and Pinheiro, 2017 [2]). Using a single recognition site in the plasmid, a nicking endonuclease and exonuclease III are added to degrade one strand of the template. A 5’-boundary oligonucleotide containing a 5’-biotinTEG modification (shown as a blue dot), a 3’-boundary oligonucleotide containing a protected 3’-end (3’-3’dT, not shown) and inner oligonucleotides are annealed onto the single stranded template. The Darwin Assembly master mix including a polymerase, ligase and dNTPs is used to extend and ligate the primers, creating the library.The assembled construct is captured via biotin-streptavidin pulldown using paramagnetic beads (shown in brown). The boundary oligonucleotides also contain PCR priming sites and Type IIs restriction sites (shown as reds overhangs) which are used for PCR recovery and subcloning into the expression backbone. The protocol is fully automated up to isothermal assembly and we are now optimizing the automation for capture, clean-up and PCR recovery steps.

## 2. Materials

Prepare all master mixes in molecular grade sterile deionised water (ddH2O) and keep them on ice. We recommend preparing the solutions immediately before use. All oligonucleotides used in these examples are provided in Table S1 and all plasmid sequences are provided in Table S2. Darwin oligo design, Darwin design files and workflows for Antha are provided as supplementary information.

### 2.1. Master mixes for Gilson Pipetmax

1. Phosphorylation master mix (5x): 1 mM Adenosine 5’-Triphosphate (ATP, New England Biolabs, NEB), 0.1 U/µL T4 Polynucleotide Kinase (T4 PNK, NEB), 1x CutSmart® buffer (NEB).
2. Single stranded plasmid master mix (3x): 3 U/µL Nb.BbvCI (NEB), 36 U/µL Exonuclease III (ExoIII, NEB), 3x CutSmart® buffer (Note 1).
3. Darwin Assembly master mix (2x): 0.05 U/µl Q5 High-Fidelity DNA polymerase (NEB), 8 U/µl Taq DNA ligase (NEB), 2 mM β-Nicotinamide adenine dinucleotide (NAD+, Sigma-Aldrich), 0.4 mM each deoxynucleotide (dNTP, Thermo Fisher Scientific), 10% (w/v) Poly(ethylene glycol) 8000 (PEG8000, Sigma-Aldrich), 2 mM 1,4-Dithiothreitol (DTT, Sigma-Aldrich) and 1x CutSmart® buffer.
4. PCR master mix: 1x KOD Xtreme Hot Start DNA Polymerase buffer (EMD Millipore), 1.2 µM (each) forward and reverse primers, 0.4 mM dNTPs, 0.4 ng/µL template DNA, 0.02 U/µL KOD Xtreme Hot Start DNA Polymerase (EMD Millipore).

### 2.2. Streptavidin Clean-up

1. Dynabeads MyOne Streptavidin C1 beads (Thermo Fisher Scientific)
2. Magnetic stand (Promega cat#: Z5342)
3. 2x BWBS-T: 20 mM Tris-HCl pH 7.4, 2 M NaCl, 0.2% v/v Tween20, 2 mM ethylenediaminetetraacetic acid (EDTA)
4. 30 mM NaOH
5. EB-T: 10 mM Tris-HCl pH 8.8, 0.1 mM EDTA, 0.01% Tween20
6. EB: 10 mM Tris-HCl pH 8.8

### 2.3. Type IIS restriction cloning and library transformation

1. SapI restriction enzyme (NEB)
2. 10x CutSmart® buffer (NEB)
3. T4 DNA ligase (NEB)
4. 1x T4 DNA ligase buffer (NEB)
5. T4 PNK (NEB)
6. T7 Express lysY/Iq (NEB)
7. NEB® 5-alpha competent *E. coli* cells (NEB)
8. 1 mM 4-(2-hydroxyethyl)-1-piperazineethanesulfonic acid (HEPES)-NaOH buffer pH 7 (Sigma-Aldrich)
9. 40 cm cell scraper (Corning)
10. Fisherbrand™ Electroporation Cuvettes Plus™ (Thermo Fisher Scientific, cat#: 15542423)
11. Corning 500 cm² Square TC-treated Culture Dish (Corning)

### 2.4. Additional Reagents and Solutions

1. UltraPure Phenol:Chloroform:Isoamyl Alcohol (25:24:1) (Thermo Fisher Scientific)
2. 4 M Ammonium acetate (Sigma-Aldrich)
3. 20 mg/mL Glycogen blue (Thermo Fisher Scientific)
4. 100% isopropanol (Sigma-Aldrich)
5. 70% ethanol [diluted from 96% Ethanol (Sigma-Aldrich) with ddH_2_O]

### 2.5 Equipment

1. Gene Pulser II electroporator (Bio-Rad Laboratories)
2. Gilson PIPETMAX™ and Antha software
3. Additional equipment and reagents for carrying out agarose gel electrophoresis
4. GeneJET PCR Purification Kit (Thermo Fisher Scientific) and GeneJET Plasmid Miniprep (Thermo Fisher Scientific) or any other commercially available PCR and plasmid purification kits.
5. KingFisher™ Duo Prime Purification System (Thermo Fisher Scientific)

## 3. Methods

The following protocol describes the automation of Darwin Assembly (DA) with biotinylated boundary oligonucleotides (see Fig. 1) specific for the pET23d_DA template and inner oligonucleotides specific for the mutagenesis of the KOD DNAP (Note 2).

### 3.1 Library template preparation

Darwin assembly imposes some constraints on the design of the initial plasmid template. Ideally, the plasmid should contain a single BbvCI site downstream of the library assembly region, binding sites for the boundary oligonucleotides used in the assembly, and no SapI restriction sites. Those modifications were introduced into the original pET23 to generate the pET23d_DA backbone used here through iPCR or QuikChange II Site-Directed Mutagenesis (Note 3), prior to KOD DNA polymerase subcloning.

1. Transform the template harbouring the gene of interest into NEB® 5-alpha competent *E. coli* cells following the recommended High Efficiency Transformation Protocol (C2987) described by commercial strain provider.
2. Isolate successful transformants and, from overnight cultures, extract the plasmid DNA to obtain at least 50 µL of a 500 ng/µL of pET23d_DA harbouring the gene of interest. Multiple methods are available for plasmid DNA isolation from bacterial cultures on this scale. We find it convenient to use multiple (5 ∼ 8) GeneJET Plasmid Miniprep (Thermo Fisher Scientific) columns and then concentrate the isolated DNA in a SpeedVac Vacuum Concentrator (Thermo Fisher Scientific) at −40°C and < 0.250 mbar for 1 h or until sufficient water has been removed from samples.

### 3.2 Library Design

Library assembly can be divided in two parts: design and construction. Library design is dependent on the application and it is not described in detail here. Nonetheless, the output of the design process are the sites to be targeted and the mutations to be introduced.

1. Design the boundary, outnest and inner mutagenic oligonucleotides. Boundary oligonucleotides form the 5’- and 3’-ends of the single stranded template during isothermal assembly (see Fig. 1), and thus anneal upstream and downstream of the region being assembled in the single stranded template. Both boundary oligonucleotides carry overhangs containing restriction sites and outnest PCR priming sites, which are described in more detail in Fig. 3.
2. Design the inner oligonucleotides, which carry the desired mutations. They should have annealing temperatures between 55°C and 60°C (assuming no mismatches) and at least 11 nucleotides on each side of the mutation, preferably until a G or C is reached.

### 3.3 Library Construction with Antha

Once the design has been generated, the process of constructing the libraries starts with entering all the components into a Darwin design file (see SI ‘Darwin Design File’) – a spreadsheet that is used by Antha as input for the experimental setup of the assembly. An overview of the custom-built Darwin Assembly element suite in Antha is shown in Fig. 2a. Each element controls distinct parts of the Darwin Assembly experiment from preparing the single-stranded template plasmid and complex mutational primer master mixes to the final preparation of mutagenesis reaction mixes for isothermal assembly, further described below.

1. Enter your experimental design in the Darwin design file (Fig. 2b and SI ‘Darwin Design File’). Define all oligonucleotide and target plasmid names, corresponding DNA sequences, DNA part stock concentrations (µM) and desired template-mutagenic primer combinations in the design file.
2. Upload this file to the ***Upload Darwin Design File*** element in the Antha cloud-based software platform.
3. Enter the experimental parameters for each part of the Darwin workflow within the appropriate elements in Antha, a complete set of suggested and validated reaction parameters as established for the original publication are provided in SI ‘Darwin Design File’.
  3.1. **Make Single Stranded DNA** – *This element generates a single stranded DNA from a double stranded plasmid by addition of a nuclease master mix.* Set the template plasmid target concentration per reaction or choose a default concentration to be used across all reactions (Fig. 2c and 2d). Define incubation time and temperature for the reaction, and for the subsequent enzyme inactivation, to trigger a prompt with appropriate instructions once the reactions have been prepared. Also specify the plate type in which to prepare the reactions (Note 4).
  3.2. **Phosphorylate Primer Mixes** – This element prepares and phosphorylates mutagenic primer mixes specified in the Darwin Design File, as well as 3’ boundary primers, by addition of a kinase master mix. Specify the primer phosphorylation reaction volume as well as mutagenic and boundary primer target concentrations to be achieved in the final reaction. It is possible to specify distinct concentrations for each mutagenic primer in a given reaction, or to set a single default concentration (Fig. 2e). Reaction incubation and enzyme inactivation times and temperatures can be set within the **Phosphorylate Primer Mixes** element to prompt the user to move reaction plates to an incubator under the desired conditions at the right time of the automation workflow. Since both **Make Single Stranded DNA** and **Phosphorylate Primer Mixes** steps are recommended to be incubated and inactivated under the same conditions, a single prompt message for the incubation and a single message for the inactivation instructions will be shown to the user once both stages have been physically executed on the liquid handling platform.
  3.3. **Isothermal Assembly** – This element mixes single-stranded template plasmid DNA, phosphorylated mutagenic and boundary primers to specified molar ratios. Following primer annealing, Darwin Assembly *master mix is added to each mixture for isothermal assembly.* Specify the total reaction volume along with single-stranded template target concentration and the molar fold excess for phosphorylated mutagenic and boundary primer mixes to single-stranded template DNA (Fig 2f). Specify an incubation time and temperature to display a prompt message containing incubation instructions, once these reactions have been physically set up.
4. Run an *in silico* simulation in Antha of the experimental workflow after having input your design file and all reaction condition parameters. During simulation, Antha will dynamically calculate the volumes of all stock reagents, given the stock concentration and desired target concentrations, required to satisfy each reaction defined in the design file (such as primer and plasmid stocks and enzymatic master mixes).
5. After the *in silico* simulation of the experimental run, open the ***Execution Details***. Antha provides microtiter plate and liquid handling platform deck layout representations for all plates pre- and post-execution (Fig. 4a). All liquid handling steps can be viewed within the ***Preview*** section (Fig. 4b), prior to physical execution of the experiment to validate the liquid handling platform will execute all mix steps as expected before needing to enter the lab.
6. ***Schedule*** the job from within the ***Execution Details Preview*** to send the liquid handling instructions to the Gilson PIPETMAX.
7. Aliquot the reaction components into the input plate as shown in the ***Execution Details Setup*** (Fig. 4a) (Note 5) and ensure all required tip racks and plates are positioned appropriately on the deck of the liquid handler as shown in the ***Execution Details Preview*** (Fig. 4b) (Note 6).
8. During the physical run, a prompt will appear once the single stranded template plasmid and phosphorylation primer mixes have been prepared. Incubate the samples as instructed on the prompt, return the samples to the deck once the incubation has completed, and press ‘continue’ to trigger subsequent liquid handling steps. Once the isothermal reactions have been prepared, a prompt with incubation instructions will appear on screen.
9. After the isothermal assembly, samples can be stored at −20°C, or used for immediate downstream processing. This is a convenient break in the assembly process and reactions are stable after 7 days (the longest storage period we have used to date).

**Figure 2.**
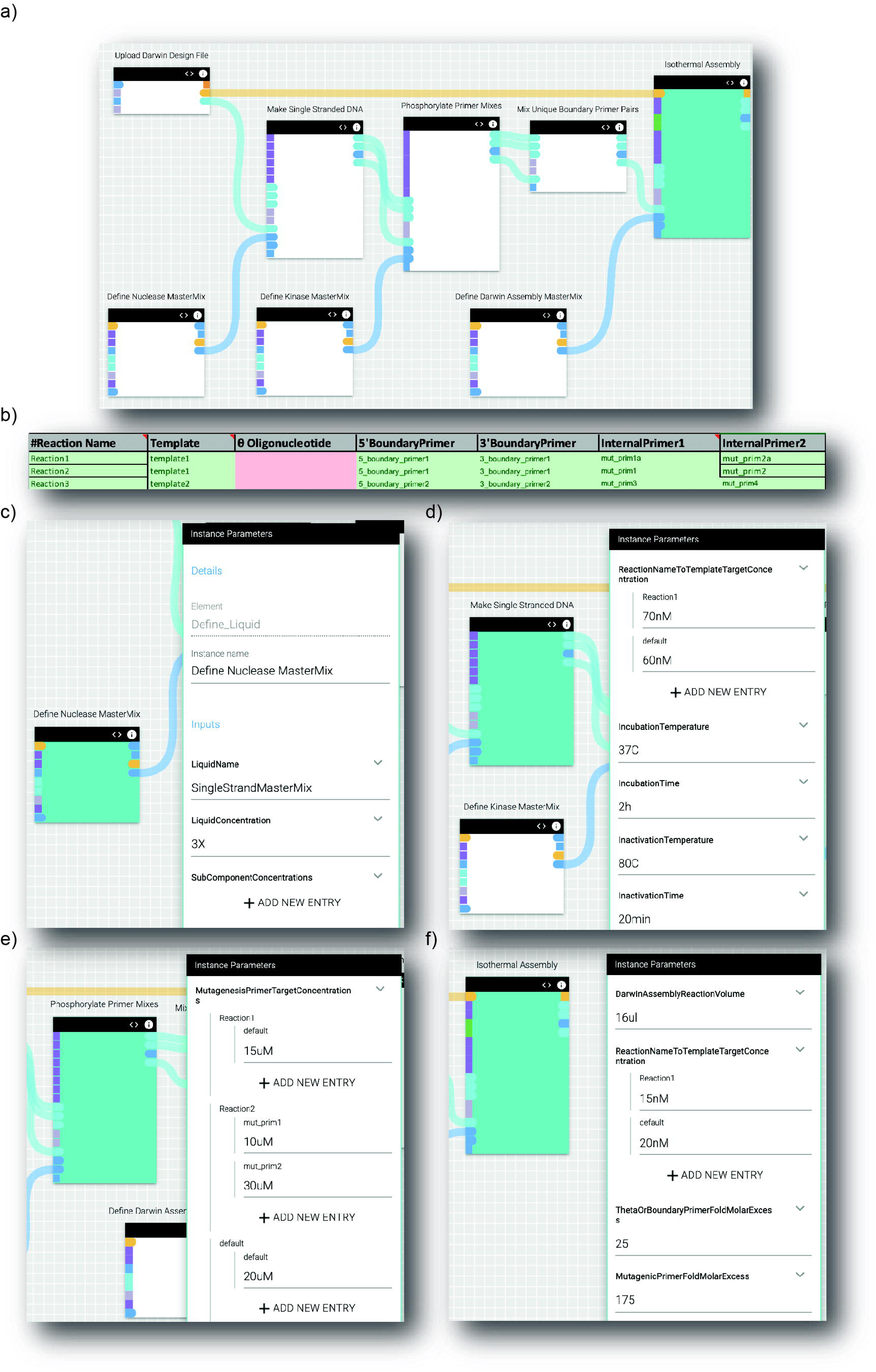
Overview of the Darwin Assembly Workflow in Antha. a)The Darwin Assembly workflow consists of multiple elements that configure distinct parts of the experiment, including generation of the single stranded plasmid containing the target gene, mixing and phosphorylation of the mutagenic and 5’-boundary primers, combining 3’- and 5’-boundary primers as required and finally combining the target DNA with the primer mixes according to specifications in the Darwin Design File. b) Example of a Darwin Assembly design file detailing template plasmids and boundary/mutagenic (internal) primers to be used in each reaction. In the above example two different plasmids (named template1 and template2) are set with a combination of mutagenic primers and appropriate boundary primers. Sequences for each of these parts are also provided in the design file (not shown). Antha will track sequences of nucleotide products throughout the workflow and perform checks for expected oligonucleotide annealing. c) Experimental parameters can be provided for each part of the workflow as shown in the ***Instance Parameter*** tab on the right, providing control and flexibility over the experiment. The ***SingleStrandMasterMix*** liquid is defined within the ***Define Liquid*** element and wired up to the ***Make Single Stranded DNA*** element. The ***Make Single Stranded DNA*** element will automatically calculate the required volumes of the ***SingleStrandMasterMix*** required to ensure a 1x final concentration in each reaction. d) Using the ***ReactionNameToTemplateTargetConcentration*** parameter within the ***Make Single Stranded DNA*** element, the final template plasmid concentration for individual reactions or a single concentration to be used for all reactions can be set. In the above example, the template plasmid concentration for Reaction1 is set to 70 nM, while 60 nM is specified for all remaining reactions (using the “default” term). e) In the ***Phosphorylate Primer Mixes*** element, the concentration of each individual mutagenic primer can be varied across reactions, if desired. In the above example, all mutagenic primers used in Reaction1 will be made up to a final concentration of 15 µM in the phosphorylation reaction, while the two mutagenic primers used in Reaction2 will be made up to 10 and 30 µM. This flexibility is particularly important when constructing libraries, where it allows controlling the expected ratio of different variants in the final population [2]. The “default” term is used to set the mutagenic primer concentration of all mutagenic primers in all remaining reactions to 20 µM. f) In the ***Isothermal Assembly element***, the desired (single-stranded) template plasmid concentration for each Darwin assembly reaction can be set. The concentration can be set specifically for a given reaction or for all reactions using the “default” term. The molar excess of boundary and mutagenic primers are also set here. The ***Isothermal Assembly*** element calculates all volumes to make up the reactions with all constituents at the specified concentrations.

**Figure 3.**
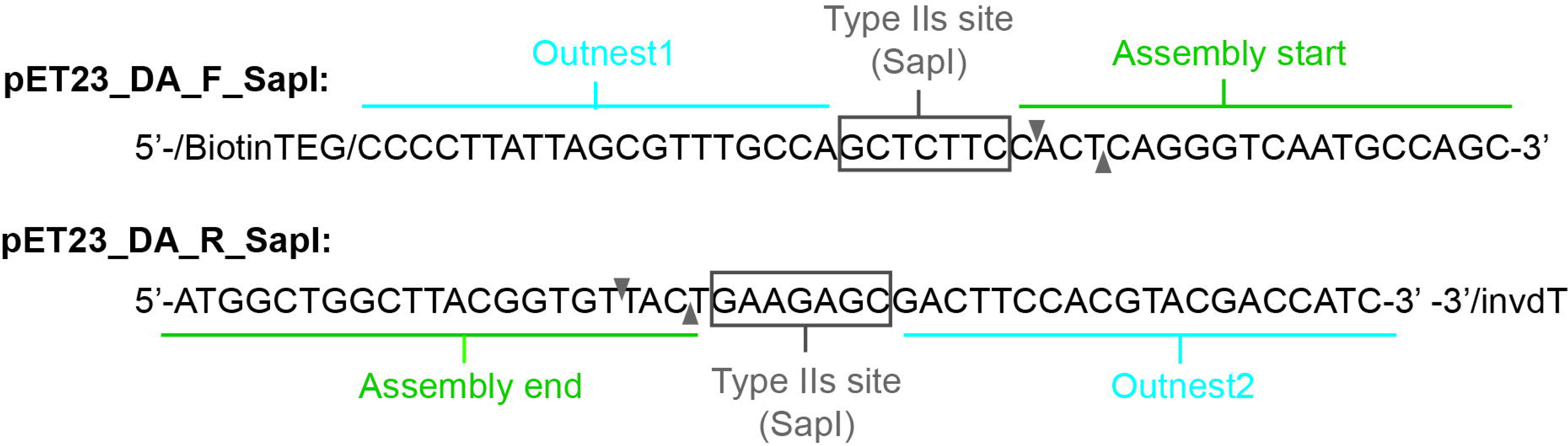
Boundary oligonucleotide design guidelines. The 5’-boundary oligonucleotide is biotinylated at its 5’-end and anneals to the start of the assembly region. It contains an overhang encoding the PCR priming site (Outnest1 - blue) and a Type IIs recognition site for Golden Gate assembly (grey). The 3’-boundary oligonucleotide binds (on the same strand) at the other end of the assembly region. Its overhang contains the reverse complement of the Type IIs recognition site and the other outnest primer (Outnest 2 - blue). The 3’-boundary oligonucleotide also contains a protected 3’-end (3’-inverted-dT) to prevent exonuclease degradation. The black triangles represent SapI cut sites in the sense and antisense strands. The strand assembled for the KOD libraries with these boundary oligonucleotides was the antisense strand.

**Figure 4.**
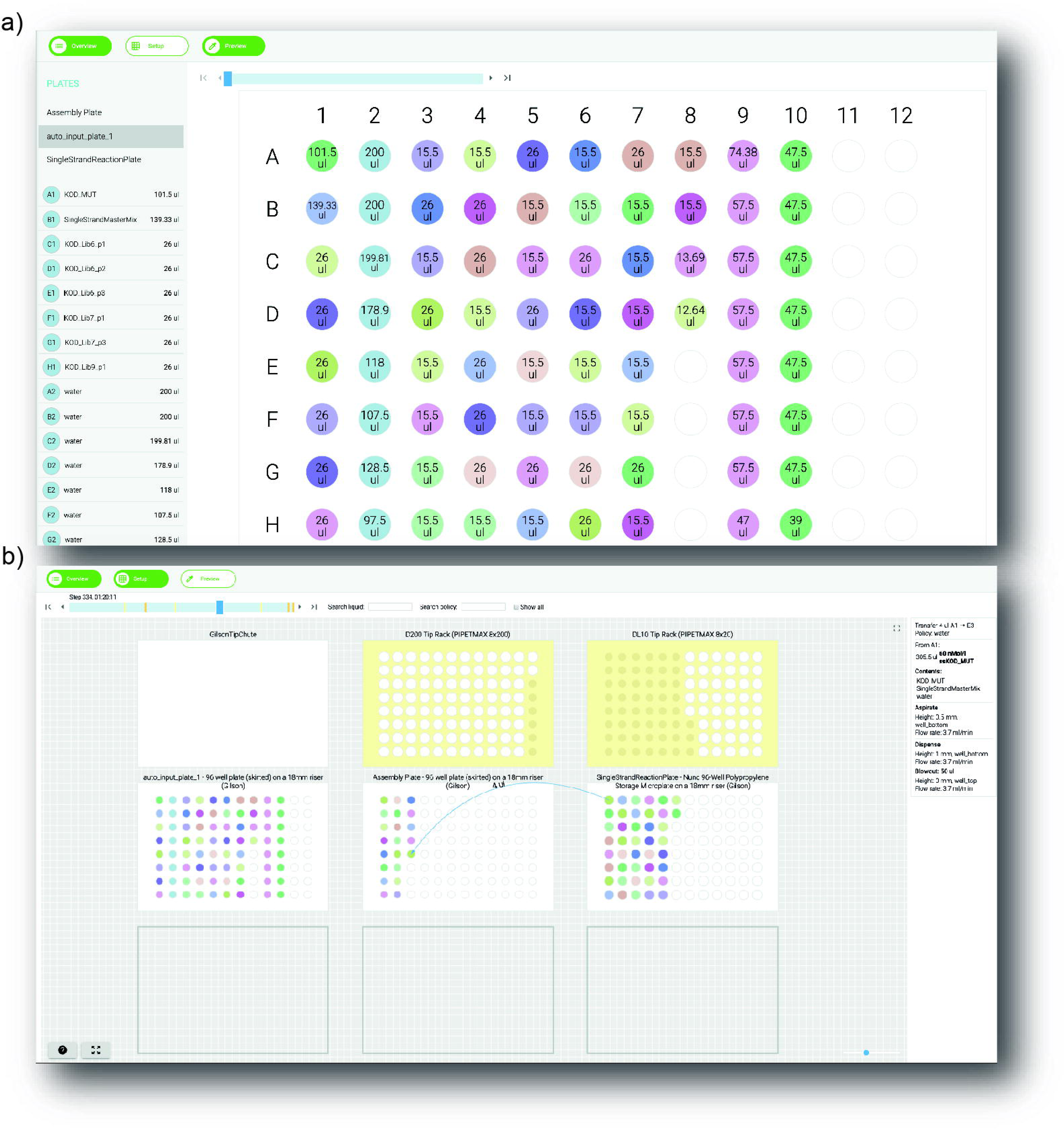
Overview of the experimental simulation results. a) The ***Execution Detail Setup*** tab visualises all experimental plates pre- and post-execution. Here, the automatically generated input file containing all liquids required to execute the experiment is shown. The volume for each liquid is calculated automatically dependent on the number of reactions assembled and the stock concentration of each reagent. b) The deck layout representation is visible in the ***Execution Detail Preview*** tab, which allows tracing of all liquid handling steps prior to any physical execution. This is controllable via the slider on the upper left, which contains further information on step number and associated timing as well as upcoming incubation prompts and tip-box changes highlighted in yellow and orange within the slider bar respectively. Low-level detail associated with each liquid handling is shown on the right-hand side.

### 3.4 Library capture (for libraries assembled with biotinylated primers)

1. Use 5 µl of Dynabeads MyOne Streptavidin C1 beads per assembly. Dilute the bead slurry in 500 µl 2x BWBS-T, capture the beads using the magnetic stand and resuspend them in fresh 500 µl 2x BWBS-T to wash them.
2. Incubate washed beads in 1 mL 2x BWBS-T for 1 h in a rotator at room temperature to block them [2]. Capture the beads using the magnetic stand and resuspend beads in 50 µl 2x BWBS-T.
3. Combine library assembly reactions (if desired) and add ddH_2_O to a total volume of 50 µl. Mix that with the 50 µl pre-blocked Dynabeads MyOne Streptavidin C1 beads.
4. Capture assembled libraries for 1 h in a rotator at room temperature.
5. Following the KingFisher™ Duo Prime Purification System protocol (Fig. 5 and SI KingFisher_DA), prepare the 96 DW plate as follows: Well B: 200 μL (37°C) 30 mM NaOH, Well C: 200 μL (37°C) 30 mM NaOH, Well D: 200 µl EB-T, Well E: 10 μL EB, Well H: KingFisher™ Duo 12 tip comb (see Fig 5).
6. Transfer 50 μL of the bead suspension to the first well (well A) of the reaction strip and start the washing program in the KingFisher™ Duo Prime Purification System.
7. Remove resuspended beads from well E and aliquot into1.5 mL Eppendorf tubes.

### 3.5 Library recovery and cloning

1. Prepare 50 µl PCRs with 1X KOD Xtreme reaction buffer, 400 µM dNTPs, 0.3 µM outnest1 and outnest2 primers (Table S1), 1 µl resuspended beads and 0.01 U/µl of KOD DNA polymerase. Incubate PCR reactions with an initial denaturation at 95°C for 2 min; 28 cycles of 98°C for 15 seconds, 67°C for 30 seconds and 68°C for 1 min 40 sec; and a final 68°C extension for 2 min (note 8).
2. Successful PCR amplification of assembled libraries can be checked by agarose gel electrophoresis (note 9).
3. Purify PCR products with GeneJET PCR Purification Kit (Thermo Fisher Scientific) following the manufacturer’s standard protocols (note 9). When multiple assemblies are to be combined into a single library, PCR products can be combined at this step and co-purified.
4. PCR amplify the pET23d_DA backbone with pET23_iPCR_DA_F1 and pET23_iPCR_DA_R1 primers (Table S1) – these primers harbour Type IIS SapI restriction sites that generate overhangs compatible with the ones generated in the library reactions. Reactions are run in 50 µl with 1X KOD Xtreme reaction buffer, 400 µM dNTPs, 0.3 µM forward and reverse primers, 10 ng of the pET23d_DA template and 0.01 U/µl of KOD DNA polymerase. Incubate PCR reactions with an initial denaturation at 95°C for 2 min; 35 cycles of 98°C for 15 seconds, 60°C for 30 seconds and 68°C for 2 min; and a final 68°C extension for 2 min.
5. Purify the backbone products with GeneJET PCR Purification Kit (Thermo Fisher Scientific).
6. Determine the approximate concentration of the purified vector fragment by UV spectrophotometry.
7. Digest 2 µg of the purified pET23d_DA per assembly with 0.4 U/μl of SapI and 0.4 U/µl of DpnI for 2 h at 37°C. DpnI is used here to selectively digest the original circular template, which is a known and expected source of background in cloning.
8. Digest 2 µg of assembled libraries with 0.4 U/µl of SapI for 2 h at 37°C.
9. Inactivate both digestions by incubating them at 65°C for 20 min as per manufacturer’s recommendation.
10. For each library assembly, mix 1 μg of digested backbone in a 1:1 molar ratio with the digested assembled library into a 100 μL ligation reaction in 1x T4 DNA ligase buffer supplemented with 40 U/μl T4 DNA ligase. Carry out the ligation overnight at room temperature.
11. Purify ligations by organic solvent extraction adding a mixture of phenol, chloroform and isoamyl alcohol (25:24:1) to the DNA in a 1:1 ratio (v/v). Vortex ligations at room temperature for 30 seconds and centrifuge ligations at 13,000 x *g* for 3 min at 4°C. Recover the aqueous phase of ligations and transfer them to a new Eppendorf tube.
12. Precipitate ligations by adding 4 M ammonium acetate in a 1:10 ratio to the DNA volume, 100% isopropanol in a 3:1 ratio to the DNA volume and 1 µl of 20 mg/mL glycogen blue to facilitate visual detection of the pellet.
13. Incubate ligations at −80°C for 5 min and centrifuge them at 13,000 x *g* for 30 min at 4°C.
14. Discard the supernatant and gently wash pellets with 300 µl of 70% isopropanol. Centrifuge them at 13,000 x *g* for another 5 min at room temperature. Discard the supernatant, dry the pellet in a heat block for 10 min at 50°C, and resuspend the pellet in 10 µl of ddH2O.
15. Grow an overnight starting culture of T7 Express lysY/Iq (NEB) in 10 mL of LB with no antibiotics at 37°C with shaking (Note 10).
16. For each microgram of DNA to be transformed, start a 200 mL fresh T7 Express lysY/Iq culture from a dilution [1% (v/v) or more dilute] of the starting overnight culture.
17. Allow the culture to reach an OD_600_ of 0.4 incubating at 37°C with shaking. Split the culture into 50 mL centrifuge tubes and pellet the cells through centrifugation at 4000 rpm (3250 x *g*) for 30 min at 4°C. Discard the supernatant and resuspend pellets in 20 mL filter-sterilised chilled 1 mM HEPES pH 7.0 buffer.
18. Combine pairs of cell suspensions in the same tube and repeat the centrifugation at 4000 rpm (3250 x *g*) for 30 min at 4°C. Discard the supernatant.
19. Repeat step 3.4.15 until a single pellet is collected (per 200 mL culture). Resuspend the pellet in 400 μl of 1 mM HEPES buffer for each ligation to be transformed.
20. Mix 400 μl of competent cells with each purified ligation and transfer it to a chilled 0.2 cm electroporation cuvette (Fisher Scientific). Transform cells with an electroporator (at 2.5 kV, 200 Ω and 25 μF).
21. Resuspend cells in 5 mL of fresh LB media and incubate them at 37°C for 1 h with shaking to allow recovery. Following incubation, centrifuge cells at 4,000 rpm (3,250 x *g*) for 5 min at 4°C, discard supernatant and resuspend the cells in 1 mL of LB.
22. Plate the 1 mL of cells in 500 cm² square TC-treated Culture Dish (Corning) containing LB agar with 50 µg/mL ampicillin (for the AmpR selection marker of pET23_DA).
23. Seal plates with plastic wrap to minimize solid media drying and incubate overnight at 37°C.
24. Harvest colonies using a 40 cm cell scraper (Corning) and resuspend them in 5 mL of storage media [LB, 50 µg/mL ampicillin and 25% (v/v) filter-sterilised glycerol] (Note 11).
25. Split resuspended cells into 5 cryovial tubes and store them at −80°C if they are to be used for directed evolution experiments.

### 3.6 Barcoding libraries - rationale

Because Darwin assembly can be used to generate very diverse libraries (> 10^7^ variants) targeting multiple distal sites, a barcoding strategy had to be developed to enable adequate sampling of those libraries. As summarised in Fig. 6, this strategy requires barcoding primers that introduce sequence diversity, anchoring primers that position the barcode adjacent to target regions and retrieval primers that generate fragments for deep sequencing.

**Figure 5.**
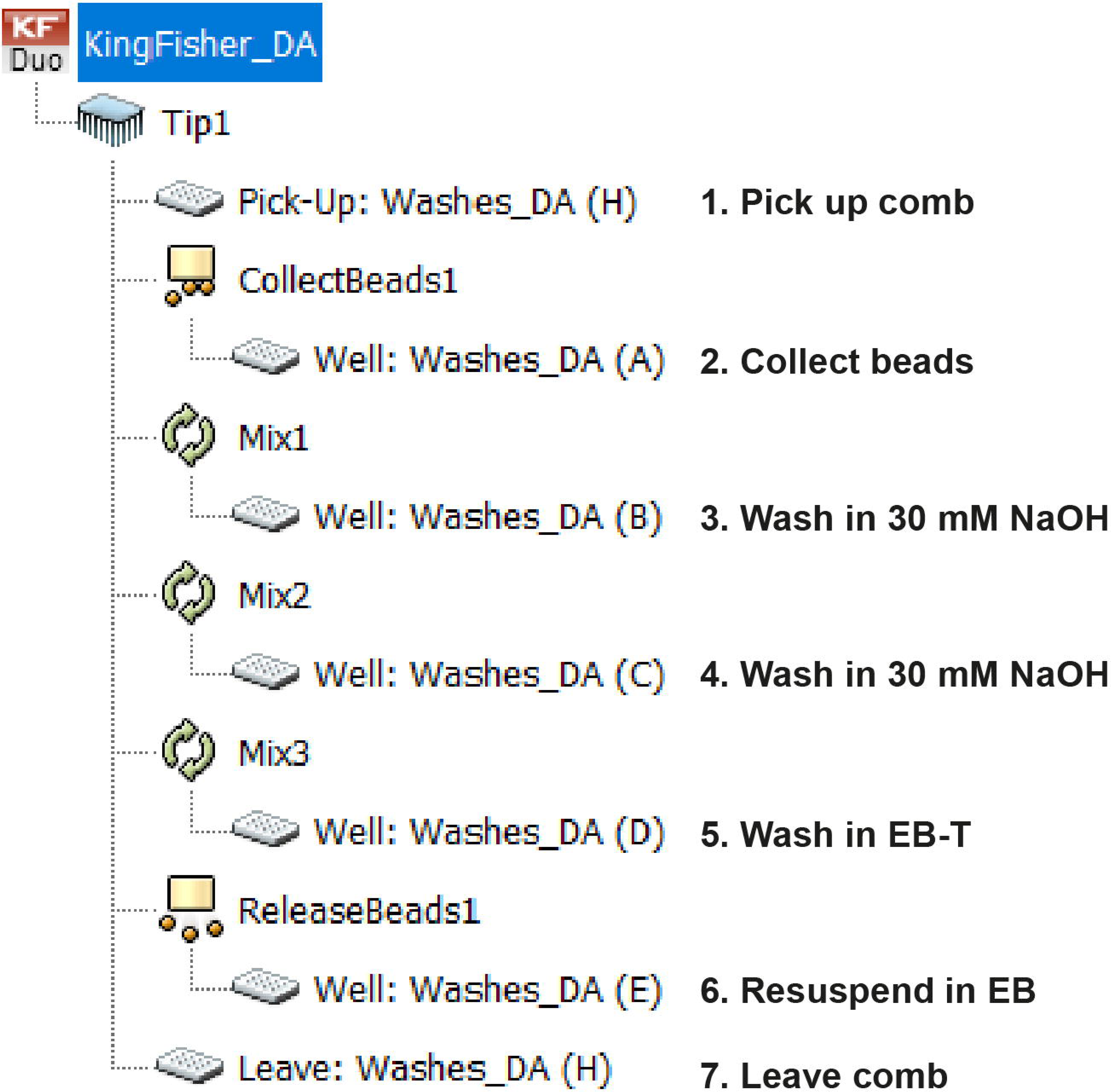
Protocol for bead washing post-capture. The washing steps were automated using the KingFisher™ Duo Prime Purification System (SI KingFisher_DA) as it allows greater parallelization and increased robustness and reproducibility. Following capture, beads are placed on well A, 200 µl of 30 mM NaOH on wells B and C, 200 µl EB-T on well D and 10 µl EB on well E. The KingFisher™ Duo 12 tip comb is picked up from well H of the 96 DW plate and proceeds to collect beads from well A with a collection time of 2 sec. Beads are transferred and mixed for 45 s at medium speed on wells B through D. The beads are finally resuspended and released in 10 µl EB (well E) following a release time of 10 s at fast speed. The KingFisher™ Duo 12 tip comb is then left on well H.

**Figure 6.**
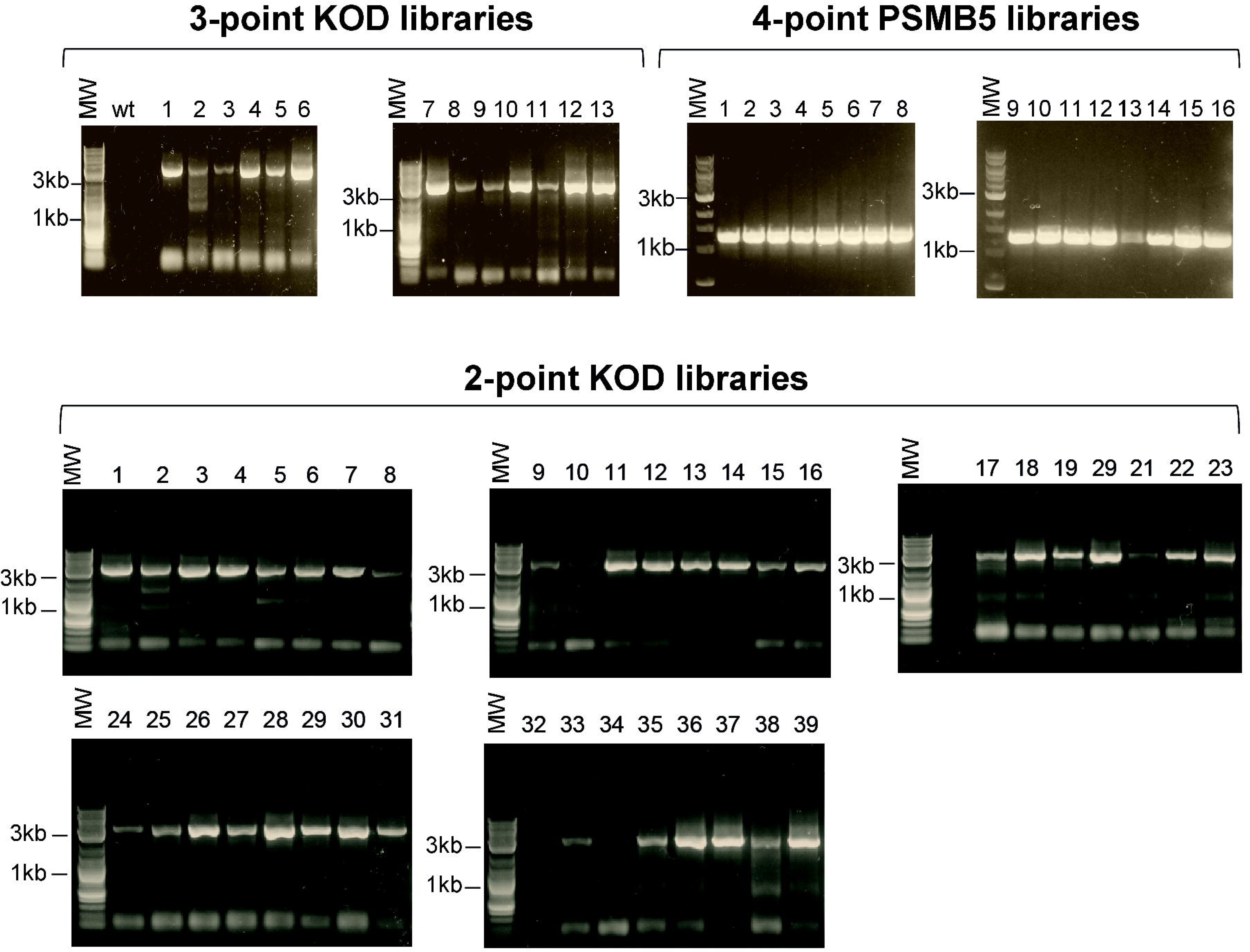
Darwin Assembly PCR recovery. PCR recovery with outnest1/outnest2 (Table S1) primers of 13 3-point site saturation mutagenesis libraries of KOD DNAP (top left gels, expected 3 kb amplicon) including a negative control with wild-type pET23_DA_KOD template. The results of 39 2-point saturation mutagenesis libraries of KOD DNAP (bottom gels, expected 3kb amplicon) are also shown. A total of 39 residues in the KOD DNAP gene were targeted. PCR recovery with outnest3/outnest4 (Table S1) of 16 4-point mutant libraries of the proteasome subunit β5 (PSMB5) gene. A total of 48 residues in the PSMB5 gene were targeted. All Darwin Assemblies were carried out using biotinylated boundary oligonucleotides using a Gilson PIPETMAX™ liquid handling platform under the control of the Antha.

**Figure 7.**
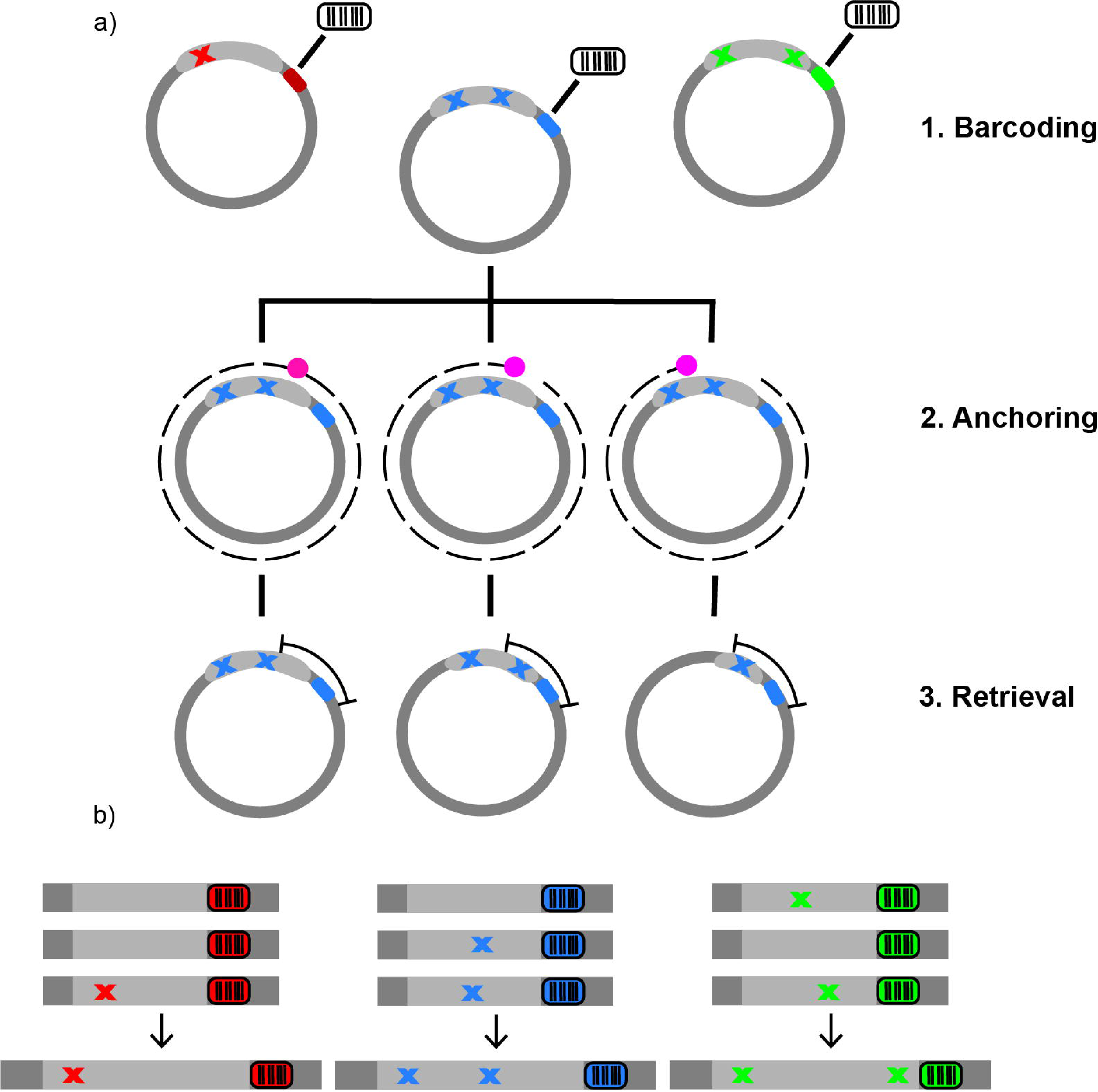
Barcoding libraries for NGS sequencing. a) PCR primers containing degeneracies (barcoding primers) are used to introduce sequence diversity outside the plasmid expression cassette creating the barcodes shown in different colors (red, blue and green) (Note 12). Primers adjacent to the barcode and to a desired gene region, termed anchoring primers, are used in iPCR to amplify the plasmid and position barcode and sequence of interest at the extremities of the molecule. A pink dot is used to emphasise the progression of the anchoring primers towards the 5’-end of the gene. Circularizing those molecules creates a PCR template in which the barcode and region of interest are juxtaposed and can be reamplified (using retrieval primers) for sequencing. b) Deep sequencing of these assembled fragments allows large libraries to be adequately sampled, as paired barcodes enable distal DNA re-assembly.

### 3.7 Barcoding libraries - Introducing barcodes to libraries

1. Split the gene into convenient 300 – 500 bp fragments that will be amplified for deep sequencing.
2. Design anchoring reverse primers against the 3′-end of each of those fragments. A single forward anchoring primer, iPCR_Part_F1 (Table S1), is needed, and it is designed against the sequence immediately upstream of the barcode in pET23d_DA. See Fig. 6 for design details.
3. Design forward retrieval primers against the 5’-end of each of the fragments to be sequenced. A single reverse retrieval primer, KOD_ amp_R1 (Table S1), is needed, and it is designed against the sequence immediately downstream of the barcode in pET23d_DA. See Fig. 6 for design details.
4. Take a small aliquot from the library glycerol stock and use it to inoculate a 10 mL culture of LB supplemented with 50 µg/mL ampicillin.
5. Incubate the culture for 3 h at 37°C with shaking.
6. Extract plasmids from the library using the GeneJET Plasmid Miniprep Kit (Thermo Fisher Scientific) following the manufacturer’s standard protocols.
7. Introduce barcodes into the library by iPCR (Note 12) in a 50 µL reaction with 1X KOD Xtreme reaction buffer, 400 µM dNTPs, 0.3 µM iPCR_Barcode_F1 and iPCR_Barcode_R1 (Table S1), 20 ng of plasmid DNA and 0.01 U/µl of KOD DNA polymerase. Incubate PCR reactions with an initial denaturation at 95°C for 2 min; 15 cycles of 98°C for 15 seconds, 60°C for 30 seconds and 68°C for 2 min 45 seconds; and a final 68°C extension for 2 min.
8. Digest PCR products in a 70 µL reaction with 0.4 U/µl of DpnI and 7 µL CutSmart® buffer.
9. Mix the digested PCR products with 40 U/μL T4 DNA ligase, 10 µL T4 DNA ligase buffer and 40 U/µL T4 polynucleotide kinase in a 100 µL reaction and incubate for 1 h at room temperature. This is a convenient point to interrupt the protocol as reactions can also proceed overnight at room temperature.
10. Purify and precipitate ligations as described in 3.5.8 – 3.5.11 and transform the barcoded library into freshly prepared competent cells as described in 3.5.12 – 3.5.22.

### 3.8 Assembling a barcoded NGS sample

1. Take a small aliquot from the library glycerol stock and use it to inoculate a 10 mL culture in LB with 50 µg/mL ampicillin.
2. Incubate the culture for 3 h at 37°C with shaking.
3. Extract plasmids from the library using the GeneJET Plasmid Miniprep Kit (Thermo Fisher Scientific).
4. Amplify 20 ng of the purified barcoded library in a 50 µl reaction with 1X KOD Xtreme reaction buffer, 400 µM dNTPs, 0.3 µM anchoring iPCR primers and 0.01 U/µL of KOD DNA polymerase. Incubate PCR reactions with an initial denaturation at 95°C for 2 min; 15 cycles of 98°C for 15 seconds, 67°C for 30 seconds and 68°C for 30 seconds/kb of the target DNA product; and a final 68°C extension for 2 min.
5. Purify PCR products with GeneJET PCR Purification Kit (Thermo Fisher Scientific).
6. Mix the digested PCR products with 40 U/μL T4 DNA ligase, 1x T4 DNA ligase buffer and 40 U/µL T4 polynucleotide kinase in a 100 µl reaction and incubate overnight at room temperature.
7. Purify and precipitate ligations as described in 3.6.8 – 3.6.11
8. Using the retrieval primers (Table S1) amplify each library fragment by PCR as described in step 3.8.4.
9. Purify the expected PCR amplicon from agarose gels stained with SYBR safe (Life Technologies) using Monarch DNA Gel Extraction kits (NEB) following manufacturer’s recommended protocol.
10. Quantify the recovered DNA using a Qubit fluorometer (Thermo Fisher Scientific).
11. Dilute or concentrate products to a final 25 µL of a 20 ng/µL to prepare samples for NGS-based amplicon sequencing (Genewiz Germany GmbH) (Note 13).
12. Using the Galaxy public server (https://usegalaxy.org) join paired-end reads on the overlapping ends allowing 0 mismatches and a minimum of 10 bp overlap using the ***FASTQ-join*** (Galaxy Version 1.1.2-484).
13. Convert FASTQ to FASTA format using ***FASTQ to FASTA converter*** (Galaxy Version 1.0.0), discarding reads with unknown bases.
14. Trim reads at the 3’- and 5’-ends using the ***Cutadapt*** (Galaxy Version 1.16.4) tool, inputting adapter sequences immediately upstream of the sequenced fragment and downstream the barcode (i.e. retrieval primer sequences) with 100% adapter overlap, no minimum or maximum length of trimmed reads and 1 bp mismatch maximum between adapter and read. Reads that did not contain adapter were discarded.
15. Export and align sequences using clustal-omega-1.2.4.
16. Using the SI python script ‘Barcoding_reconstruction’, input the name of your MSA files in FASTA format and a sequence immediately before the barcode under ‘Splitting at barcode.’
17. Run the script. The reconstructed gene sequences based on matching barcodes should be saved as a text file, which can be then used to identify the multiple mutations introduced in a single gene (see Fig. 8).

**Figure 8.**
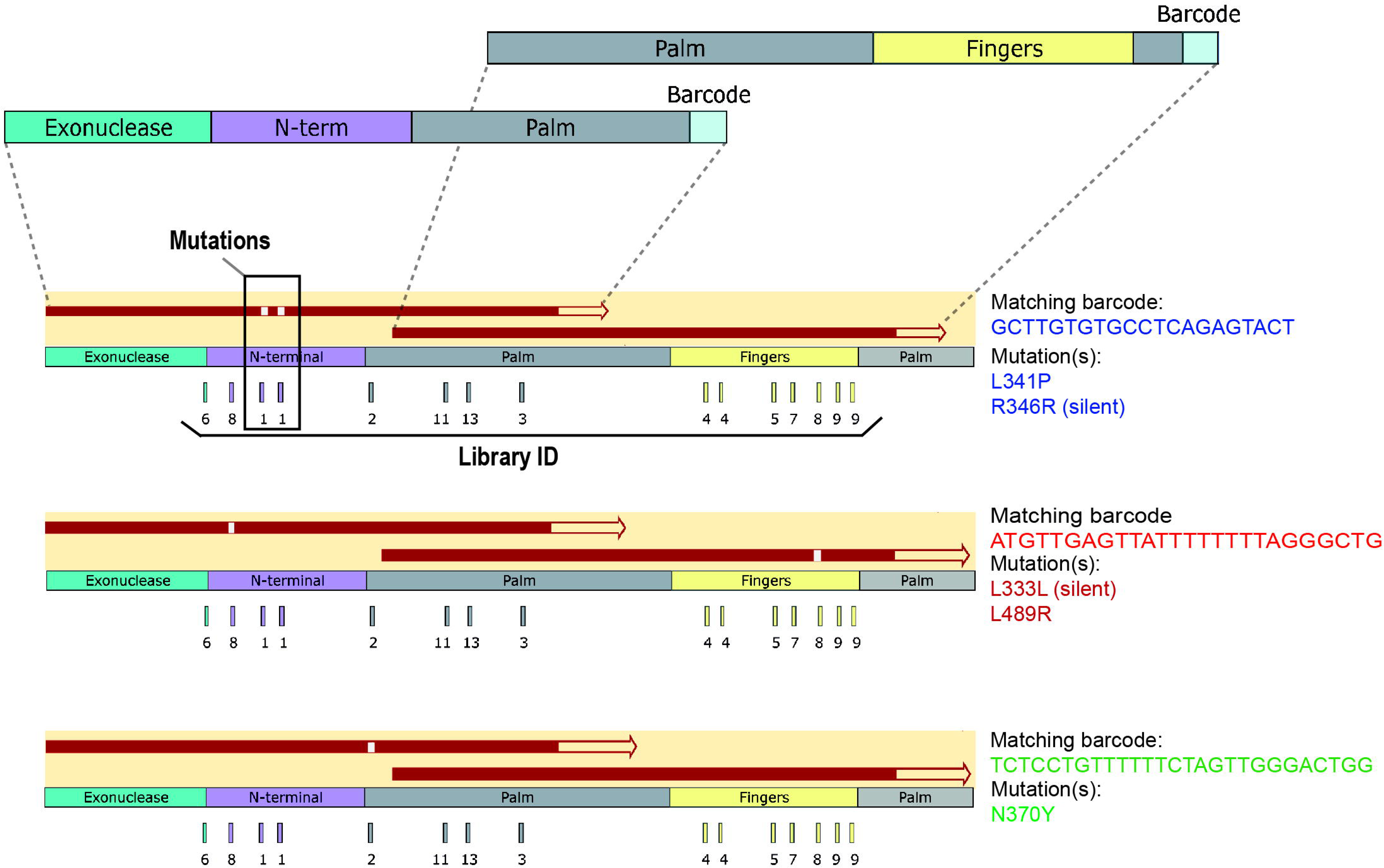
Reconstruction of entire read lengths using barcodes. Two barcoded fragments covering different regions of the gene were generated from the 3-point mutant library of KOD DNAP. Amplicons were commercially sequenced (Genewiz Germany GmbH). KOD DNAP subdomains are shown in different colours: exonuclease (teal), N-terminal (purple), palm (grey) and fingers (yellow). The barcode is shown in light blue. Fragments were paired based on the barcode to reconstruct the entire gene. Three pairs of fragments sharing the same barcode are shown here, with each read aligned to the reference KOD DNAP sequence. In all cases, we can see that mutations are localized to targeted residues and within the coded diversity (NNK).

### 4. Notes

1. The recommended enzyme concentration for single stranded plasmid generation is template concentration dependant. Quantities were estimated for the digestion of 19 µg of template, necessary for 14 assemblies. Typically 9U/µg DNA of Nb.BbvCI, 120 U/µg DNA of ExoIII and 3x CutSmart® buffer are used for the generation of the 3x master mix.
2. Darwin Assembly requires inner mutagenic oligonucleotides, biotinylated boundary oligonucleotides or a single theta boundary oligonucleotide, and outnest primers. Ensure the outnest primers share no complementarity to the wild-type template.
3. The pET23d_DA template has been optimised by introducing the DA fragment, containing a BbvCI nicking site as well as an optimised boundary oligonucleotide priming site and a barcoding priming site, between the 3’-end of the gene and the T7 terminator sequence by iPCR. The pET23d_DA template also contains a single NotI restriction site in its plasmid replication origin introduced by iPCR, which can be used for enzymatic degradation during DA with the theta oligonucleotide. Alternatively, the gene can be subcloned into pET29_DA, which, as shown here, was successfully nicked with BspQI. BtsI nicking endonuclease has also been successfully tested with pET29_DA vector. Alternative boundary oligonucleotide priming sites can be introduced to prevent non-specific binding if the insert gene shares partial complementarity. If other templates are to be used, introduce a nicking endonuclease site (to date BbvCI, BspQI and BtsI have been tested successfully in DA) onto the strand that will be degraded and remove additional sites on the opposite strand if necessary. We have found that placing the nicking endonuclease site downstream of the assembly region yields better results than placing it upstream.
4. Output plates and risers can be adjusted according to the liquid handler requirements, compatible thermocyclers available and reaction volume required. One 96-well reaction plate can be used to hold phosphorylation primer mixes and the reactions for creating single stranded template-plasmids. This can be achieved by connecting two output parameters from the ***Make Single Stranded DNA*** element (***PlateNameUsed*** and ***WellsUsed*** parameters) with input parameters on the ***Phosphorylate Primer Mixes*** element (***NewPlateName*** and ***AvoidWells***, respectively) (Fig. 2a).
5. Antha determines the location of reagents in the input plate. However, to avoid having to aliquot several primers into specific locations in a 96-well plate, a csv file specifying the desired plate layout can be uploaded into Antha. Since several incubations will be carried out, enzyme mixes can be aliquoted into the input plate just before the relevant reaction takes place. A PCR-cooler on-deck can also be used.
6. If the workflow requires multiple tip boxes and these need to be changed during the execution, the protocol will pause and prompt the user to change tip boxes as required. The estimated timing of these events can be viewed in the ***Execution Details Preview*** slide bar where they are highlighted in orange (Fig. 4).
7. Post assembly PCR can be carried out with any DNA polymerase, but higher fidelity enzymes are recommended for longer assemblies. We have successfully used MyTaq (Bioline), Q5 Hot Start DNA polymerase (NEB) and KOD Xtreme (Merck) in this step. Nonetheless, we find that, in general, KOD Xtreme robustly delivers better PCR yields.
8. Thermocycling parameters can be adjusted according to the DNA polymerase being used, the melting temperature (T_m_) of the outnest primers, the length of the assembly product and the synthesis rate of the DNA polymerase.
9. If multiple amplicons are observed during agarose gel electrophoresis, PCR parameters can be optimized or the expected PCR amplicon purified from the agarose gel using a Monarch DNA Gel Extraction Kit (New England Biolabs).
10. Libraries were transformed in T7 Express lysY/Iq (NEB), since the constructs used here were under the regulation of a T7 promoter. Other expression strains and plasmids can be used.
11. If the library is not going to be directly expressed from glycerol stocks, the plasmids can be extracted by resuspending all the colonies in 10 mL of LB. Centrifuge cells and discard the supernatant. Measure the pellet wet weight and extract plasmids by alkaline lysis with SDS [Miniprep Alkaline lysis with SDS], adjusting the volumes according to the pellet wet weight in comparison to a 2 mL overnight culture which should weight approximately 6 mg.
12. Barcoding primers presented here are compatible with the pET23d_DA but if an alternative backbone is used, design barcoding iPCR primers by introducing degeneracies in the oligos. Our approach of introducing (NNK)_7_ as the barcode yields an expected 3×10^10^ unique variants. Barcodes can be introduced during or after cloning the assembled library.
13. Other deep sequencing platforms can be used according to amplicon length and library size.

## Supporting information

Scripts and SI Tables

## Acknowledgement

VBP thanks BBSRC (grant BB/N01023X/1 - invivoXNA), ERC (grant 336936 – HNAepisome) and the Rega Institute (grant ZL57031000). VBP and PH thank FWO (grant G0H7618N Odysseus).

## Conflict of Interest

MK is an employee of Synthace, a for-profit company developing Antha.

