## Supplementary material for "Antha-guided Automation of Darwin Assembly for the Construction of Bespoke Gene Libraries": Scripts and SI Tables: Barcoding_reconstruction.py

```
# -*- coding: utf-8 -*-
"""
@author: PH
"""

from Bio.Seq import Seq
from Bio import SeqIO
import numpy as np
from Bio import AlignIO
import pandas as pd  

#import MSA datasets

A = AlignIO.read("KOD_fragment1_MSA", "fasta")
B = AlignIO.read("KOD_fragment2_MSA", "fasta")

#to string
A_df = pd.DataFrame(A)   
B_df = pd.DataFrame(B)  

A_str = A_df[A_df.columns[:]].apply(
    lambda x: ''.join(x.dropna().astype(str)),
    axis=1
)

B_str = B_df[B_df.columns[:]].apply(
    lambda x: ''.join(x.dropna().astype(str)),
    axis=1
)

#splitting at barcode
A_split = A_str.str.split(pat ='TGAGGTAGGAT', expand =True)
B_split = B_str.str.split(pat ='TGAGGTAGGAT', expand =True)

A_bar = A_split[2]
B_bar = B_split[2]


#remove InDels
A_bar = A_bar.str.split(pat ='-', expand =True)
B_bar = B_bar.str.split(pat ='-', expand =True)

A_bar = A_bar[A_bar.columns[:]].apply(
    lambda x: ''.join(x.dropna().astype(str)),
    axis=1
)

B_bar = B_bar[B_bar.columns[:]].apply(
    lambda x: ''.join(x.dropna().astype(str)),
    axis=1
)


#match barcodes
A_df['Bar'] = A_bar 
B_df['Bar'] = B_bar

#keeping unique sequences
A_subset= A_df.drop_duplicates(subset=['Bar'], keep='last')
B_subset= B_df.drop_duplicates(subset=['Bar'], keep='last')

AB_rec = pd.merge(A_subset, B_subset, left_on='Bar', right_on='Bar', how='left').drop('Bar', axis=1)

#generate MSA
AB_rec['MSA'] = AB_rec[AB_rec.columns[:]].apply(
    lambda x: ''.join(x.dropna().astype(str)),
    axis=1
)

rec = AB_rec['MSA'].str.split(pat ='-', expand =True)
rec = rec[rec.columns[:]].apply(
    lambda x: ''.join(x.dropna().astype(str)),
    axis=1
)

#removing un-paired fragments
msa_df = rec['MSA'].to_frame()
list_seqs = msa_df["MSA"].values
list_names = msa_df.index.values
list_seq = list_seqs.astype(str)
list_name = list_names.astype(str)

#df to fasta

ofile = open("my_fastav3.txt", "w")

for i in range(len(list_seqs)):

    ofile.write(">" + list_name[i] + "\n" +list_seq[i] + "\n")

ofile.close()
```
