## Supplementary material for "Antha-guided Automation of Darwin Assembly for the Construction of Bespoke Gene Libraries": Scripts and SI Tables: Table S1.docx

### Table S1: Oligonucleotides used for Darwin Assembly and Barcoding Libraries

| **Darwin Assembly on pET23_DA_KOD** | | |
| --- | --- | --- |
| **Category** | **Name** | **Sequence** |
| **Biotinylated** | pET23_DA_F_SapI | /5BiotinTEG/CCCCTTATTAGCGTTTGCCAGCTCTTCCACTCAGGGTCAATGCCAGC |
|  | pET23_DA_R_SapI | ATGGCTGGCTTACGGTGTTACTGAAGAGCGACTTCCACGTACGACCATC/3InvdT/ |
| **Outnest** | Outnest1 | CCCCTTATTAGCGTTTGCCA |
|  | Outnest2 | GATGGTCGTACGTGGAAGTC |
| **Backbone amplification** | pET23_iPCR_DA_R1 | AAAGCTCTTCCAGTGATTTTTCTCTGGTCC |
|  | pET23_iPCR_DA_F1 | AAAGCTCTTCCTACTGCTGAGGAATGAGCC |
| **Inner Mutagenic** | KOD_Lib1_p1 | GGGACGGATACACNNKGATCTCTATCCTGTG |
|  | KOD_Lib1_split_2 | CGGCCAGTCCNNKTGGGACGTCTC |
|  | KOD_Lib1_p2n3 | CGGCCAGTCCNNKTGGGACGTCTCCNNKTCCAGCACTGG |
|  | KOD_Lib1_Split_3 | GGGACGTCTCCNNKTCCAGCACTGGC |
|  | KOD_Lib2_p1n2 | GGGACAAGATANNKGAGCATCCAGCAGTTATTNNKATCTACGAGTAC |
|  | KOD_Lib2_p3 | GAGCTGGCCCCGNNKAAGCCCGATG |
|  | KOD_Lib2_Split_1 | GGGACAAGATANNKGAGCATCCAGCAG |
|  | KOD_Lib2_Split_2 | CAGCAGTTATTNNKATCTACGAGTAC |
|  | KOD_Lib3_p1n2 | GATCCCTGTACNNKNNKATCATCATCACCCACAAC |
|  | KOD_Lib3_p3 | CTaGGAGACCTCNNKGAGGAGAGGCAG |
|  | KOD_Lib3_Split_1 | GATCCCTGTACNNKTCAATCATCATCACCC |
|  | KOD_Lib3_Split_2 | CCCTGTACCCCNNKATCATCATCACCC |
|  | KOD_Lib4_p1 | CGATGGAGGCCNNKCTTTCTCGCTTAATC |
|  | KOD_Lib4_p2n3 | GACCTCCTAGAGNNKAGGCAGAAGNNKAAGAAGAAGATG |
|  | KOD_Lib4_Split_2 | GACCTCCTAGAGNNKAGGCAGAAGATAAAG |
|  | KOD_Lib4_Split_3 | GAGAGGCAGAAGNNKAAGAAGAAGATG |
|  | KOD_Lib5_p1n2 | CCGTTTATGAANNKNNKTTCGGTCAGCCG |
|  | KOD_Lib5_p3 | CGATCGAGAGGNNKCTCCTCGATTAC |
|  | KOD_Lib5_Split_1 | GCCGTTTATGAANNKGTCTTCGGTCAG |
|  | KOD_Lib5_Split_2 | GTTTATGAAGCCNNKTTCGGTCAGCC |
|  | KOD_Lib6_p1 | GATAAGACGGACGNNKAACCTGCCCAC |
|  | KOD_Lib6_p2 | CTTGGGAAGGAGNNKCTTCCGATGGAG |
|  | KOD_Lib6_p3 | GAAGCTCCTCGATNNKAGGCAGAGGttg |
|  | KOD_Lib7_p1 | GTGATAAGACGGNNKATAAACCTGCCC |
|  | KOD_Lib7_p2 | GAGTTCCTTCCGNNKGAGGCCCAGC |
|  | KOD_Lib7_p3 | CTCCTCGATTACNNKCAGAGGttgATC |
|  | KOD_Lib8_p1 | CTGTGATAAGANNKACGATAAACCTG |
|  | KOD_Lib8_p2 | GATGGAGGCCCAGNNKTCTCGCTTAATC |
|  | KOD_Lib8_p3 | GttgATCAAGATCNNKGCAAACAGCTAC |
|  | KOD_Lib9_p1 | GACGGCAGAGCNNKGAAGGAGGCTATG |
|  | KOD_Lib9_p2n3 | CAAACAGCTACNNKGGTTACTACNNKTATGCAAGGGC |
|  | KOD_Lib9_Split_2 | CAAACAGCTACNNKGGTTACTACGG |
|  | KOD_Lib9_Split_3 | CTACGGTTACTACNNKTATGCAAGGGC |
|  | KOD_Lib10_p1 | GTAAAAGAGCCCGAGNNKGGGTTGTGGGAG |
|  | KOD_Lib10_p2n3 | GCGGCTTCTTCNNKNNKAAGAAGAAGTATG |
|  | KOD_Lib10_Split_2 | GCGGCTTCTTCNNKACGAAGAAGAAGTATG |
|  | KOD_Lib10_Split_3 | GCGGCTTCTTCGTCNNKAAGAAGAAGTATG |
|  | KOD_Lib11_p1n2 | GGAGGCTATGTANNKNNKCCCGAGAGAGGG |
|  | KOD_Lib11_p3 | CTTCTTCGTCACGNNKAAGAAGTATGC |
|  | KOD_Lib11_Split_1 | GGAGGCTATGTANNKGAGCCCGAGAG |
|  | KOD_Lib11_Split_2 | GCTATGTAAAANNKCCCGAGAGAGG |
|  | KOD_Lib12_p1n2 | CTATCCTGTGATANNKCGGACGATAAACCTGNNKACATACACGCTTG |
|  | KOD_Lib12_split_2 | CGATAAACCTGNNKACATACACGCTTG |
|  | KOD_Lib12_p3 | GAGATTGTGAGGNNKGACTGGAGCGAG |
|  | KOD_Lib12_Split_1 | CTATCCTGTGATANNKCGGACGATAAAC |
|  | KOD_Lib13_p1 | CCGAGAGAGGGNNKTGGGAGAACATAG |
|  | KOD_Lib13_p3 | GAATTCTGAGAGCCNNKGGTTACCGCAAG |
|  | KOD_Lib13_p2 | CTTGAAGCTTTGNNKAAGGACGGTGAC |
| **Darwin Assembly on pET23_DA_KOD** | | |
| **Category** | **Name** | **Sequence** |
| **Biotinylated** | pET29_DA_F_BsaI | /5BioTinTEG/GGCAACTAGAAGGCACAGTCGGTCTCCTTCCTTTCGGGCTTTGTTAGCAGC |
|  | pET29_DA_R_BsaI | GTTAAACAAAATTATTTCTAGAGGGGAATTGTTATCCGCGAGACCCAGTGGGTTCTCTAGTTAGCC/3InvdT/ |
| **Outnest** | Outnest3 | GGCAACTAGAAGGCACAGTC |
|  | Outnest4 | GGCTAACTAGAGAACCCACTG |
| **Backbone amplification** | pET29_iPCR_DA_R1 | GAGTCAGGTCTCAGGAAGCTGAGTTGGCTGCTG |
|  | pET29_iPCR_DA_F1 | GAGTCAGGTCTCATCCGCTCACAATTCCCCTATAGTGAG |
| **Inner Mutagenic** | 001_PSMB5_G70 | CTTCAAGTTCCGCCATNNKGTCATAGTTGCAGCTGACTCCAGGGCTA |
|  | 001_PSMB5_V71 | CTTCAAGTTCCGCCATGGANNKATAGTTGCAGCTGACTCCAGGGCTA |
|  | 001_PSMB5_I72 | CTTCAAGTTCCGCCATGGAGTCNNKGTTGCAGCTGACTCCAGGGCTA |
|  | 001_PSMB5_V73 | CAAGTTCCGCCATGGAGTCATANNKGCAGCTGACTCCAGGGCTACAGCGG |
|  | 001_PSMB5_A74 | CAAGTTCCGCCATGGAGTCATAGTTNNKGCTGACTCCAGGGCTACAGCGG |
|  | 001_PSMB5_A75 | CAAGTTCCGCCATGGAGTCATAGTTGCANNKGACTCCAGGGCTACAGCGG |
|  | 001_PSMB5_D76 | CCATGGAGTCATAGTTGCAGCTNNKTCCAGGGCTACAGCGGGTGCTTACAT |
|  | 001_PSMB5_S77 | CCATGGAGTCATAGTTGCAGCTGACNNKAGGGCTACAGCGGGTGCTTACAT |
|  | 001_PSMB5_R78 | CCATGGAGTCATAGTTGCAGCTGACTCCNNKGCTACAGCGGGTGCTTACAT |
|  | 001_PSMB5_A79 | CATAGTTGCAGCTGACTCCAGGNNKACAGCGGGTGCTTACATTGCCTCCCAG |
|  | 001_PSMB5_T80 | CATAGTTGCAGCTGACTCCAGGGCTNNKGCGGGTGCTTACATTGCCTCCCAG |
|  | 001_PSMB5_A81 | CATAGTTGCAGCTGACTCCAGGGCTACANNKGGTGCTTACATTGCCTCCCAG |
|  | 001_PSMB5_G82 | TGACTCCAGGGCTACAGCGNNKGCTTACATTGCCTCCCAGACGGTGA |
|  | 001_PSMB5_A83 | TGACTCCAGGGCTACAGCGGGTNNKTACATTGCCTCCCAGACGGTGA |
|  | 001_PSMB5_Y84 | TGACTCCAGGGCTACAGCGGGTGCTNNKATTGCCTCCCAGACGGTGA |
|  | 001_PSMB5_I85 | GGCTACAGCGGGTGCTTACNNKGCCTCCCAGACGGTGAAGAAGGTGATA |
|  | 001_PSMB5_A86 | GGCTACAGCGGGTGCTTACATTNNKTCCCAGACGGTGAAGAAGGTGATA |
|  | 001_PSMB5_S87 | GGCTACAGCGGGTGCTTACATTGCCNNKCAGACGGTGAAGAAGGTGATA |
|  | 001_PSMB5_Q88 | CGGGTGCTTACATTGCCTCCNNKACGGTGAAGAAGGTGATAGAGATCAACCCA |
|  | 001_PSMB5_T89 | CGGGTGCTTACATTGCCTCCCAGNNKGTGAAGAAGGTGATAGAGATCAACCCA |
|  | 001_PSMB5_V90 | CGGGTGCTTACATTGCCTCCCAGACGNNKAAGAAGGTGATAGAGATCAACCCA |
|  | 001_PSMB5_K91 | TTGCCTCCCAGACGGTGNNKAAGGTGATAGAGATCAACCCATACCTGCTA |
|  | 001_PSMB5_K92 | TTGCCTCCCAGACGGTGAAGNNKGTGATAGAGATCAACCCATACCTGCTA |
|  | 001_PSMB5_V93 | TTGCCTCCCAGACGGTGAAGAAGNNKATAGAGATCAACCCATACCTGCTA |
|  | 001_PSMB5_I94 | CCAGACGGTGAAGAAGGTGNNKGAGATCAACCCATACCTGCTAGGCAC |
|  | 001_PSMB5_E95 | CCAGACGGTGAAGAAGGTGATANNKATCAACCCATACCTGCTAGGCAC |
|  | 001_PSMB5_I96 | CCAGACGGTGAAGAAGGTGATAGAGNNKAACCCATACCTGCTAGGCAC |
|  | 001_PSMB5_N97 | CGGTGAAGAAGGTGATAGAGATCNNKCCATACCTGCTAGGCACCATGGCT |
|  | 001_PSMB5_P98 | CGGTGAAGAAGGTGATAGAGATCAACNNKTACCTGCTAGGCACCATGGCT |
|  | 001_PSMB5_Y99 | CGGTGAAGAAGGTGATAGAGATCAACCCANNKCTGCTAGGCACCATGGCT |
|  | 001_PSMB5_L100 | AGGTGATAGAGATCAACCCATACNNKCTAGGCACCATGGCTGGGGGC |
|  | 001_PSMB5_L101 | AGGTGATAGAGATCAACCCATACCTGNNKGGCACCATGGCTGGGGGC |
|  | 001_PSMB5_G102 | AGGTGATAGAGATCAACCCATACCTGCTANNKACCATGGCTGGGGGC |
|  | 001_PSMB5_T103 | CAACCCATACCTGCTAGGCNNKATGGCTGGGGGCGCAGCGG |
|  | 001_PSMB5_M104 | CAACCCATACCTGCTAGGCACCNNKGCTGGGGGCGCAGCGG |
|  | 001_PSMB5_A105 | CAACCCATACCTGCTAGGCACCATGNNKGGGGGCGCAGCGG |
|  | 001_PSMB5_G106 | GCTAGGCACCATGGCTNNKGGCGCAGCGGATTGCAGCTTCTGG |
|  | 001_PSMB5_G107 | GCTAGGCACCATGGCTGGGNNKGCAGCGGATTGCAGCTTCTGG |
|  | 001_PSMB5_A108 | GCTAGGCACCATGGCTGGGGGCNNKGCGGATTGCAGCTTCTGG |
|  | 001_PSMB5_A109 | GGCTGGGGGCGCANNKGATTGCAGCTTCTGGGAACGGCTG |
|  | 001_PSMB5_D110 | GGCTGGGGGCGCAGCGNNKTGCAGCTTCTGGGAACGGCTG |
|  | 001_PSMB5_C111 | GGCTGGGGGCGCAGCGGATNNKAGCTTCTGGGAACGGCTG |
|  | 001_PSMB5_S112 | GGCGCAGCGGATTGCNNKTTCTGGGAACGGCTGTTGGCTCG |
|  | 001_PSMB5_F113 | GGCGCAGCGGATTGCAGCNNKTGGGAACGGCTGTTGGCTCG |
|  | 001_PSMB5_W114 | GGCGCAGCGGATTGCAGCTTCNNKGAACGGCTGTTGGCTCG |
|  | 001_PSMB5_E115 | GCGGATTGCAGCTTCTGGNNKCGGCTGTTGGCTCGGCAATGTCGA |
|  | 001_PSMB5_R116 | GCGGATTGCAGCTTCTGGGAANNKCTGTTGGCTCGGCAATGTCGA |
|  | 001_PSMB5_L117 | GCGGATTGCAGCTTCTGGGAACGGNNKTTGGCTCGGCAATGTCGA |
|  | 001_PSMB5_E8 | CTTGCCAGCGTGTTGGAAAGACCGCTACCGGTGAAC |
| **Barcoding pET23_DA_KOD Libraries for NGS** | | |
| **Category** | **Name** | **Sequence** |
| **Barcoding** | iPCR_Barcode_F1 | NNKNNKNNKNNKNNKNNKNNKGGCTGGCTTACGG |
|  | iPCR_Barcode_R1 | ATCCTACCTCATCTCGGAGC |
| **Anchoring** | iPCR_Part_F1 | GCTCCGAGATGAGGTAGGAT |
|  | iPCR_Part_KOD_R1 | CGAGACGTTGTGGGTGATG |
|  | iPCR_Part_KOD_R2 | CACTCCTTGCAGTACCAGC |
| **Retrieval** | KOD_amp_R1 | CAGTAACACCGTAAGCCAGC |
|  | KOD_amp_F1 | GGTCAGCCGAAGGAGAAG |
|  | KOD_amp_F2 | GAGCTGGCCAGAAGACG |
