## Supplementary material for "Antha-guided Automation of Darwin Assembly for the Construction of Bespoke Gene Libraries": Scripts and SI Tables: Table S2.docx

### Table S2: Plasmid Sequences

**pET23_DA**

tggcgaatgggacgcgccctgtagcggcgcattaagcgcggccgctgtggtggttacgcgcagcgtgaccgctacacttgccagcgccctagcgcccgctcctttcgctttcttcccttcctttctcgccacgttcgccggctttccccgtcaagctctaaatcgggggctccctttagggttccgatttagtgctttacggcacctcgaccccaaaaaacttgattagggtgatggttcacgtagtgggccatcgccctgatagacggtttttcgccctttgacgttggagtccacgttctttaatagtggactcttgttccaaactggaacaacactcaaccctatctcggtctattcttttgatttataagggattttgccgatttcggcctattggttaaaaaatgagctgatttaacaaaaatttaacgcgaattttaacaaaatattaacgtttacaatttcaggtggcacttttcggggaaatgtgcgcggaacccctatttgtttatttttctaaatacattcaaatatgtatccgctcatgagacaataaccctgataaatgcttcaataatattgaaaaaggaagagtatgagtattcaacatttccgtgtcgcccttattcccttttttgcggcattttgccttcctgtttttgctcacccagaaacgctggtgaaagtaaaagatgctgaagatcagttgggtgcacgagtgggttacatcgaactggatctcaacagcggtaagatccttgagagttttcgccccgaagaacgttttccaatgatgagcacttttaaagttctgctatgtggcgcggtattatcccgtattgacgccgggcaagagcaactcggtcgccgcatacactattctcagaatgacttggttgagtactcaccagtcacagaaaagcatcttacggatggcatgacagtaagagaattatgcagtgctgccataaccatgagtgataacactgcggccaacttacttctgacaacgatcggaggaccgaaggagctaaccgcttttttgcacaacatgggggatcatgtaactcgccttgatcgttgggaaccggagctgaatgaagccataccaaacgacgagcgtgacaccacgatgcctgcagcaatggcaacaacgttgcgcaaactattaactggcgaactacttactctagcttcccggcaacaattaatagactggatggaggcggataaagttgcaggaccacttctgcgctcggcccttccggctggctggtttattgctgataaatctggagccggtgagcgtgggtctcgcggtatcattgcagcactggggccagatggtaagccctcccgtatcgtagttatctacacgacggggagtcaggcaactatggatgaacgaaatagacagatcgctgagataggtgcctcactgattaagcattggtaactgtcagaccaagtttactcatatatactttagattgatttaaaacttcatttttaatttaaaaggatctaggtgaagatcctttttgataatctcatgaccaaaatcccttaacgtgagttttcgttccactgagcgtcagaccccgtagaaaagatcaaaggatcttcttgagatcctttttttctgcgcgtaatctgctgcttgcaaacaaaaaaaccaccgctaccagcggtggtttgtttgccggatcaagagctaccaactctttttccgaaggtaactggcttcagcagagcgcagataccaaatactgtacttctagtgtagccgtagttaggccaccacttcaagaactctgtagcaccgcctacatacctcgctctgctaatcctgttaccagtggctgctgccagtggcgataagtcgtgtcttaccgggttggactcaagacgatagttaccggataaggcgcagcggtcgggctgaacggggggttcgtgcacacagcccagcttggagcgaacgacctacaccgaactgagatacctacagcgtgagctatgagaaagcgccacgcttcccgaagggagaaaggcggacaggtatccggtaagcggcagggtcggaacaggagagcgcacgagggagcttccagggggaaacgcctggtatctttatagtcctgtcgggtttcgccacctctgacttgagcgtcgatttttgtgatgctcgtcaggggggcggagcctatggaaaaacgccagcaacgcggcctttttacggttcctggccttttgctgcgttatcccctgattctgtggccttttgcgcgtctgcgttatcccctgattctgatgttctttcctgcgttatcccctgattctgtggataaccgtattaccgcctttgagtgagctgcgttatcccctgattctgctgataccgctcgccgcagccgaacgaccgagcgcagcgagtcagtgagcgaggaagcggaatatcgcctgatgcggtattttctccttacgcatctgtgcggtatttcacaccgcaatggtgcactctcagtacaatctgctctgatgccgcatagttaagccagtatacactccgctatcgctacgtgactgggtcatggctgcgccccgacacccgccaacacccgctgacgcgccctgacgggcttgtctgctcccggcatccgcttacagacaagctgtgaccgtctccgggagctgcatgtgtcagaggttttcaccgtcatcaccgaaacgcgcgaggcagctgcggtaaagctcatcagcgtggtcgtgaagcgattcacagatgtctgcctgttcatccgcgtccagctcgttgagtttctccagaagcgttaatgtctggcttctgataaagcgggccatgttaagggcggttttttcctgtttggtcactgatgcctccgtgtaagggggatttctgttcatgggggtaatgataccgatgaaacgagagaggatgctcacgatacgggttactgatgatgaacatgcccggttactggaacgttgtgagggtaaacaactggcggtatggatgcggcgggaccagagaaaaatcactcagggtcaatgccagcgcttcgttaatacagatgtaggtgttccacagggtagccagcagcatcctgcgatgcagatccggaacataatggtgcagggcgctgacttccgcgtttccagactttacgaaacacggaaaccgaagaccattcatgttgttgctcaggtcgcagacgttttgcagcagcagtcgcttcacgttcgctcgcgtatcggtgattcattctgctaaccagtaaggcaaccccgccagcctagccgggtcctcaacgacaggagcacgatcatgcgcacccgtggccaggacccaacgctgcccgagatctcgatcccgcgaaattaatacgactcactatagggagaccacaacggtttccctctagaaataattttgtttaactttaagaaggagatataccatggatcctctagagtcgacctgcaggcatgcaagcttgcggccacacaggagatagtcatacatgaaacacaaagaggagaaattaactgatccggctgctaacaaagcccgaaaggaagctgagttggctgctgccaccgctgagcaataactagcataaccccttggggcctctaaacgggtcttgaggggttttttgctgaaaggaggaactatatccggat

**pET23_DA_KOD**

tggcgaatgggacgcgccctgtagcggcgcattaagcgcggccgctgtggtggttacgcgcagcgtgaccgctacacttgccagcgccctagcgcccgctcctttcgctttcttcccttcctttctcgccacgttcgccggctttccccgtcaagctctaaatcgggggctccctttagggttccgatttagtgctttacggcacctcgaccccaaaaaacttgattagggtgatggttcacgtagtgggccatcgccctgatagacggtttttcgccctttgacgttggagtccacgttctttaatagtggactcttgttccaaactggaacaacactcaaccctatctcggtctattcttttgatttataagggattttgccgatttcggcctattggttaaaaaatgagctgatttaacaaaaatttaacgcgaattttaacaaaatattaacgtttacaatttcaggtggcacttttcggggaaatgtgcgcggaacccctatttgtttatttttctaaatacattcaaatatgtatccgctcatgagacaataaccctgataaatgcttcaataatattgaaaaaggaagagtatgagtattcaacatttccgtgtcgcccttattcccttttttgcggcattttgccttcctgtttttgctcacccagaaacgctggtgaaagtaaaagatgctgaagatcagttgggtgcacgagtgggttacatcgaactggatctcaacagcggtaagatccttgagagttttcgccccgaagaacgttttccaatgatgagcacttttaaagttctgctatgtggcgcggtattatcccgtattgacgccgggcaagagcaactcggtcgccgcatacactattctcagaatgacttggttgagtactcaccagtcacagaaaagcatcttacggatggcatgacagtaagagaattatgcagtgctgccataaccatgagtgataacactgcggccaacttacttctgacaacgatcggaggaccgaaggagctaaccgcttttttgcacaacatgggggatcatgtaactcgccttgatcgttgggaaccggagctgaatgaagccataccaaacgacgagcgtgacaccacgatgcctgcagcaatggcaacaacgttgcgcaaactattaactggcgaactacttactctagcttcccggcaacaattaatagactggatggaggcggataaagttgcaggaccacttctgcgctcggcccttccggctggctggtttattgctgataaatctggagccggtgagcgtgggtctcgcggtatcattgcagcactggggccagatggtaagccctcccgtatcgtagttatctacacgacggggagtcaggcaactatggatgaacgaaatagacagatcgctgagataggtgcctcactgattaagcattggtaactgtcagaccaagtttactcatatatactttagattgatttaaaacttcatttttaatttaaaaggatctaggtgaagatcctttttgataatctcatgaccaaaatcccttaacgtgagttttcgttccactgagcgtcagaccccgtagaaaagatcaaaggatcttcttgagatcctttttttctgcgcgtaatctgctgcttgcaaacaaaaaaaccaccgctaccagcggtggtttgtttgccggatcaagagctaccaactctttttccgaaggtaactggcttcagcagagcgcagataccaaatactgtacttctagtgtagccgtagttaggccaccacttcaagaactctgtagcaccgcctacatacctcgctctgctaatcctgttaccagtggctgctgccagtggcgataagtcgtgtcttaccgggttggactcaagacgatagttaccggataaggcgcagcggtcgggctgaacggggggttcgtgcacacagcccagcttggagcgaacgacctacaccgaactgagatacctacagcgtgagctatgagaaagcgccacgcttcccgaagggagaaaggcggacaggtatccggtaagcggcagggtcggaacaggagagcgcacgagggagcttccagggggaaacgcctggtatctttatagtcctgtcgggtttcgccacctctgacttgagcgtcgatttttgtgatgctcgtcaggggggcggagcctatggaaaaacgccagcaacgcggcctttttacggttcctggccttttgctgcgttatcccctgattctgtggccttttgcgcgtctgcgttatcccctgattctgatgttctttcctgcgttatcccctgattctgtggataaccgtattaccgcctttgagtgagctgcgttatcccctgattctgctgataccgctcgccgcagccgaacgaccgagcgcagcgagtcagtgagcgaggaagcggaatatcgcctgatgcggtattttctccttacgcatctgtgcggtatttcacaccgcaatggtgcactctcagtacaatctgctctgatgccgcatagttaagccagtatacactccgctatcgctacgtgactgggtcatggctgcgccccgacacccgccaacacccgctgacgcgccctgacgggcttgtctgctcccggcatccgcttacagacaagctgtgaccgtctccgggagctgcatgtgtcagaggttttcaccgtcatcaccgaaacgcgcgaggcagctgcggtaaagctcatcagcgtggtcgtgaagcgattcacagatgtctgcctgttcatccgcgtccagctcgttgagtttctccagaagcgttaatgtctggcttctgataaagcgggccatgttaagggcggttttttcctgtttggtcactgatgcctccgtgtaagggggatttctgttcatgggggtaatgataccgatgaaacgagagaggatgctcacgatacgggttactgatgatgaacatgcccggttactggaacgttgtgagggtaaacaactggcggtatggatgcggcgggaccagagaaaaatcactcagggtcaatgccagcgcttcgttaatacagatgtaggtgttccacagggtagccagcagcatcctgcgatgcagatccggaacataatggtgcagggcgctgacttccgcgtttccagactttacgaaacacggaaaccgaagaccattcatgttgttgctcaggtcgcagacgttttgcagcagcagtcgcttcacgttcgctcgcgtatcggtgattcattctgctaaccagtaaggcaaccccgccagcctagccgggtcctcaacgacaggagcacgatcatgcgcacccgtggccaggacccaacgctgcccgagatctcgatcccgcgaaattaatacgactcactatagggagaccacaacggtttccctctagaaataattttgtttaactttaagaaggagatataccatggatcctctagagtcgacctgcaggcatgcaagcttgcggccacacaggagatagtcatacatgaaacacaaagaggagaaattaactatgagaggatctcaccatcaccatcaccatacggatccaagcggcctggtgccgcgcggcagcatgatcctcgacactgactacataaccgaggatggaaagcctgtcataagaattttcaagaaggaaaacggcgagtttaagattgagtacgaccggacttttgaaccctacttctacgccctcctgaaggacgattctgccattgaggaagtcaagaagataaccgccgagaggcacgggacggttgtaacggttaagcgggttgaaaaggttcagaagaagttcctagggagaccagttgaggtctggaaactctactttactcatccgcaggacgaaccagcgataagggacaagatacgagagcatccagcagttattgacatctacgagtacgacatacccttcgccaagcgctacctcatagacaagggattagtgccaatggaaggcgacgaggagctgaaaatgctcgccttcgcgattgcgactctctaccatgagggcgaggagttcgccgaggggccaatccttatgataagctacgccgacgaggaaggggccagggtgataacttggaagaacgtggatctcccctacgttgacgtcgtctcgacggagagggagatgataaagcgcttcctccgtgttgtgaaggagaaagacccggacgttctcataacctacaacggcgacaacttcgacttcgcctatctgaaaaagcgctgtgaaaagctcggaataaacttcgccctcggaagggatggaagcgagccgaagattcagaggatgggcgacaggtttgccgtcgaagtgaagggacggatacacttcgatctctatcctgtgataagacggacgataaacctgcccacatacacgcttgaggccgtttatgaagccgtcttcggtcagccgaaggagaaggtttacgctgaggaaataaccacagcctgggaaaccggcgagaaccttgagagagtcgcccgctactcgatggaagatgcgaaggtcacatacgagcttgggaaggagttccttccgatggaggcccagctttctcgcttaatcggccagtccctctgggacgtctcccgctccagcactggcaacctcgttgagtggttcctcctcaggaaggcctatgagaggaatgagctggccccgaacaagcccgatgaaaaggagctggccagaagacggcagagctatgaaggaggctatgtaaaagagcccgagagagggttgtgggagaacatagtgtacctagattttagatccctgtacccctcaatcatcatcacccacaacgtctcgccggatacgctcaacagagaaggatgcaaggaatatgacgttgccccacaggtcggccaccgcttctgcaaggacttcccaggatttatcccgagcctgctaggagacctcctagaggagaggcagaagataaagaagaagatgaaggccacgattgacccgatcgagaggaagctcctcgattacaggcagaggttgatcaagatcctggcaaacagctactacggttactacggctatgcaagggcgcgctggtactgcaaggagtgtgcagagagcgtaacggcctggggaagggagtacataacgatgaccatcaaggagatagaggaaaagtacggctttaaggtaatctacagcgacaccgacggattttttgccacaatacctggagccgatgctgaaaccgtcaaaaagaaggctatggagttcctcaagtatatcaacgccaaacttccgggcgcgcttgagctcgagtacgagggcttctacaaacgcggcttcttcgtcacgaagaagaagtatgcggtgatagacgaggaaggcaagataacaacgcgcggacttgagattgtgaggcgtgactggagcgagatagcgaaagagacgcaggcgagggttcttgaagctttgctaaaggacggtgacgtcgagaaggccgtgaggatagtcaaagaagttaccgaaaagctgagcaagtacgaggttccgccggagaagctggtgatccacgagcagataacgagggatttaaaggactacaaggcaaccggtccccacgttgccgttgccaagaggttggccgcgagaggagtcaaaatacgccctggaacggtgataagctacatcgtgctcaagggctctgggaggataggcgacagggcgataccgttcgacgagttcgacccgacgaagcacaagtacgacgccgagtactacattgagaaccaggttctcccagccgttgagagaattctgagagccttcggttaccgcaaggaagacctgcgctaccagaagacgagacaggttggtttgagtgcttggctgaagccgaagggaacttgatcgatgctccgagatgaggtaggatggctggcttacggtgttactgctgaggaatgagccatcctcgagcaccaccaccaccaccactgagatccggctgctaacaaagcccgaaaggaagctgagttggctgctgccaccgctgagcaataactagcataaccccttggggcctctaaacgggtcttgaggggttttttgctgaaaggaggaactatatccggat

**pET29a_DA**

atccggatatagttcctcctttcagcaaaaaacccctcaagacccgtttagaggccccaaggggttatgctagttattgctcagcggtggcagcagccaactcagcttcctttcgggctttgttagcagccggatctcagtggtggtggtggtggtgctcgagtgcggccgcaagcttgtcgacggagctcgaattcggatccgatatcagccatggaaccgcgtggcaccagggtacccagatctgggctgtccatgtgctggcgttcgaatttagcagcagcggtttctttcatatgtatatctccttcttaaagttaaacaaaattatttctagaggggaattgttatccgctcacaattcccctatagtgagtcgtattaatttcgcgggatcgagatcgatctcgatcctctacgccggacgcatcgtggccggcatcaccggcgccacaggtgcggttgctggcgcctatatcgccgacatcaccgatggggaagatcgggctcgccacttcgggctcatgagcgcttgtttcggcgtgggtatggtggcaggccccgtggccgggggactgttgggcgccatctccttgcatgcaccattccttgcggcggcggtgctcaacggcctcaacctactactgggctgcttcctaatgcaggagtcgcataagggagagcgtcgagatcccggacaccatcgaatggcgcaaaacctttcgcggtatggcatgatagcgcccggaagagagtcaattcagggtggtgaatgtgaaaccagtaacgttatacgatgtcgcagagtatgccggtgtctcttatcagaccgtttcccgcgtggtgaaccaggccagccacgtttctgcgaaaacgcgggaaaaagtggaagcggcgatggcggagctgaattacattcccaaccgcgtggcacaacaactggcgggcaaacagtcgttgctgattggcgttgccacctccagtctggccctgcacgcgccgtcgcaaattgtcgcggcgattaaatctcgcgccgatcaactgggtgccagcgtggtggtgtcgatggtagaacgaagcggcgtcgaagcctgtaaagcggcggtgcacaatcttctcgcgcaacgcgtcagtgggctgatcattaactatccgctggatgaccaggatgccattgctgtggaagctgcctgcactaatgttccggcgttatttcttgatgtctctgaccagacacccatcaacagtattattttctcccatgaagacggtacgcgactgggcgtggagcatctggtcgcattgggtcaccagcaaatcgcgctgttagcgggcccattaagttctgtctcggcgcgtctgcgtctggctggctggcataaatatctcactcgcaatcaaattcagccgatagcggaacgggaaggcgactggagtgccatgtccggttttcaacaaaccatgcaaatgctgaatgagggcatcgttcccactgcgatgctggttgccaacgatcagatggcgctgggcgcaatgcgcgccattaccgagtccgggctgcgcgttggtgcggacatctcggtagtgggatacgacgataccgaagacagctcatgttatatcccgccgttaaccaccatcaaacaggattttcgcctgctggggcaaaccagcgtggaccgcttgctgcaactctctcagggccaggcggtgaagggcaatcagctgttgcccgtctcactggtgaaaagaaaaaccaccctggcgcccaatacgcaaaccgcctctccccgcgcgttggccgattcattaatgcagctggcacgacaggtttcccgactggaaagcgggcagtgagcgcaacgcaattaatgtaagttagctcactcattaggcaccgggatctcgaccgatgcccttgagagccttcaacccagtcagctccttccggtgggcgcggggcatgactatcgtcgccgcacttatgactgtcttctttatcatgcaactcgtaggacaggtgccggcagcgctctgggtcattttcggcgaggaccgctttcgctggagcgcgacgatgatcggcctgtcgcttgcggtattcggaatcttgcacgccctcgctcaagccttcgtcactggtcccgccaccaaacgtttcggcgagaagcaggccattatcgccggcatggcggccccacgggtgcgcatgatcgtgctcctgtcgttgaggacccggctaggctggcggggttgccttactggttagcagaatgaatcaccgatacgcgagcgaacgtgaagcgactgctgctgcaaaacgtctgcgacctgagcaacaacatgaatggtcttcggtttccgtgtttcgtaaagtctggaaacgcggaagtcagcgccctgcaccattatgttccggatctgcatcgcaggatgctgctggctaccctgtggaacacctacatctgtattaacgaagcgctggcattgaccctgagtgatttttctctggtcccgccgcatccataccgccagttgtttaccctcacaacgttccagtaaccgggcatgttcatcatcagtaacccgtatcgtgagcatcctctctcgtttcatcggtatcattacccccatgaacagaaatcccccttacacggaggcatcagtgaccaaacaggaaaaaaccgcccttaacatggcccgctttatcagaagccagacattaacgcttctggagaaactcaacgagctggacgcggatgaacaggcagacatctgtgaatcgcttcacgaccacgctgatgagctttaccgcagctgcctcgcgcgtttcggtgatgacggtgaaaacctctgacacatgcagctcccggagacggtcacagcttgtctgtaagcggatgccgggagcagacaagcccgtcagggcgcgtcagcgggtgttggcgggtgtcggggcgcagccatgacccagtcacgtagcgatagcggagtgtatactggcttaactatgcggcatcagagcagattgtactgagagtgcaccatatatgcggtgtgaaataccgcacagatgcgtaaggagaaaataccgcatcaggcgctcttccgcttcctcgctcactgactcgctgcgctcggtcgttcggctgcggcgagcggtatcagctcactcaaaggcggtaatacggttatccacagaatcaggggataacgcaggaaagaacatgtgagcaaaaggccagcaaaaggccaggaaccgtaaaaaggccgcgttgctggcgtttttccataggctccgcccccctgacgagcatcacaaaaatcgacgctcaagtcagaggtggcgaaacccgacaggactataaagataccaggcgtttccccctggaagctccctcgtgcgctctcctgttccgaccctgccgcttaccggatacctgtccgcctttctcccttcgggaagcgtggcgctttctcatagctcacgctgtaggtatctcagttcggtgtaggtcgttcgctccaagctgggctgtgtgcacgaaccccccgttcagcccgaccgctgcgccttatccggtaactatcgtcttgagtccaacccggtaagacacgacttatcgccactggcagcagccactggtaacaggattagcagagcgaggtatgtaggcggtgctacagagttcttgaagtggtggcctaactacggctacactagaaggacagtatttggtatctgcgctctgctgaagccagttaccttcggaaaaagagttggtagctcttgatccggcaaacaaaccaccgctggtagcggtggtttttttgtttgcaagcagcagattacgcgcagaaaaaaaggatctcaagaagatcctttgatcttttctacggggtctgacgctcagtggaacgaaaactcacgttaagggattttggtcatgaacaataaaactgtctgcttacataaacagtaatacaaggggtgttatgagccatattcaacgggaaacgtcttgctctaggccgcgattaaattccaacatggatgctgatttatatgggtataaatgggctcgcgataatgtcgggcaatcaggtgcgacaatctatcgattgtatgggaagcccgatgcgccagagttgtttctgaaacatggcaaaggtagcgttgccaatgatgttacagatgagatggtcagactaaactggctgacggaatttatgcctcttccgaccatcaagcattttatccgtactcctgatgatgcatggttactcaccactgcgatccccgggaaaacagcattccaggtattagaagaatatcctgattcaggtgaaaatattgttgatgcgctggcagtgttcctgcgccggttgcattcgattcctgtttgtaattgtccttttaacagcgatcgcgtatttcgtctcgctcaggcgcaatcacgaatgaataacggtttggttgatgcgagtgattttgatgacgagcgtaatggctggcctgttgaacaagtctggaaagaaatgcataaacttttgccattctcaccggattcagtcgtcactcatggtgatttctcacttgataaccttatttttgacgaggggaaattaataggttgtattgatgttggacgagtcggaatcgcagaccgataccaggatcttgccatcctatggaactgcctcggtgagttttctccttcattacagaaacggctttttcaaaaatatggtattgataatcctgatatgaataaattgcagtttcatttgatgctcgatgagtttttctaagaattaattcatgagcggatacatatttgaatgtatttagaaaaataaacaaataggggttccgcgcacatttccccgaaaagtgccacctgaaattgtaaacgttaatattttgttaaaattcgcgttaaatttttgttaaatcagctcattttttaaccaataggccgaaatcggcaaaatcccttataaatcaaaagaatagaccgagatagggttgagtgttgttccagtttggaacaagagtccactattaaagaacgtggactccaacgtcaaagggcgaaaaaccgtctatcagggcgatggcccactacgtgaaccatcaccctaatcaagttttttggggtcgaggtgccgtaaagcactaaatcggaaccctaaagggagcccccgatttagagcttgacggggaaagccggcgaacgtggcgagaaaggaagggaagaaagcgaaaggagcgggcgctagggcgctggcaagtgtagcggtcacgctgcgcgtaaccaccacacccgccgcgcttaatgcgccgctacagggcgcgtcccattcgcca

**pET29a_DA_PSMB5**

atccggatatagttcctcctttcagcaaaaaacccctcaagacccgtttagaggccccaaggggttatgctagttattgctcagcggtggcagcagccaactcagcttcctttcgggctttgttagcagccggatctcagtggtggtggtggtggtgctcgagtgcggccgcaagcttgcaacgggtttgccgccagaacacaggaccggttctagagcgctgccaccatggcgcttgccagcgtgttggagagaccgctaccggtgaaccagcgcgggtttttcggacttgggggtcgtgcagatctgctggatctaggtccagggagtctcagtgatggtctgagcctggccgcgccaggctggggtgtcccagaGgagccGggaatcgaaatgcttcatggaacaaccacccttgccttcaagttccgccatggagtcatagttgcagctgactccagggctacagcgggtgcttacattgcctcccagacggtgaagaaggtgatagagatcaacccatacctgctaggcaccatggctgggggcgcagcggattgcagcttctgggaacggctgttggctcggcaatgtcgaatctatgagcttcgaaataaggaacgcatctctgtagcagctgcctccaaactgcttgccaacatggtgtatcagtacaaaggcatggggctgtccatgggcaccatgatctgtggctgggataagagaggccctggcctctactacgtggacagtgaagggaaccggatttcaggggccaccttctctgtaggttctggctctgtgtatgcatatggggtcatggatcggggctattcctatgacctggaagtggagcaggcctatgatctggcccgtcgagccatctaccaagccacctacagagatgcctactcaggaggtgcagtcaacctctaccacgtgcgggaggatggctggatccgagtctccagtgacaatgtggctgatctacatgagaagtatagtggctctacccccgattacaaagacgatgacgataagggatccggcgcaacaaacttctctctgctgaaacaagccggagatgtcgaagagaatcctggaccgaccgagtacaagcccacggtgcgcctcgccacccgcgacgacgtccccagggccgtacgcaccctcgccgccgcgttcgccgacatcagccatggaaccgcgtggcaccagggtacccagatctgggctgtccatgtgctggcgttcgaatttagcagcagcggtttctttcatatgtatatctccttcttaaagttaaacaaaattatttctagaggggaattgttatccgctcacaattcccctatagtgagtcgtattaatttcgcgggatcgagatcgatctcgatcctctacgccggacgcatcgtggccggcatcaccggcgccacaggtgcggttgctggcgcctatatcgccgacatcaccgatggggaagaccgggctcgccacttcgggctcatgagcgcttgtttcggcgtgggtatggtggcaggccccgtggccgggggactgttgggcgccatctccttgcatgcaccattccttgcggcggcggtgctcaacggcctcaacctactactgggctgcttcctaatgcaggagtcgcataagggagagcgtcgagatcccggacaccatcgaatggcgcaaaacctttcgcggtatggcatgatagcgcccggaagagagtcaattcagggtggtgaatgtgaaaccagtaacgttatacgatgtcgcagagtatgccggtgtctcttatcagaccgtttcccgcgtggtgaaccaggccagccacgtttctgcgaaaacgcgggaaaaagtggaagcggcgatggcggagctgaattacattcccaaccgcgtggcacaacaactggcgggcaaacagtcgttgctgattggcgttgccacctccagtctggccctgcacgcgccgtcgcaaattgtcgcggcgattaaatctcgcgccgatcaactgggtgccagcgtggtggtgtcgatggtagaacgaagcggcgtcgaagcctgtaaagcggcggtgcacaatcttctcgcgcaacgcgtcagtgggctgatcattaactatccgctggatgaccaggatgccattgctgtggaagctgcctgcactaatgttccggcgttatttcttgatgtctctgaccagacacccatcaacagtattattttctcccatgaagacggtacgcgactgggcgtggagcatctggtcgcattgggtcaccagcaaatcgcgctgttagcgggcccattaagttctgtctcggcgcgtctgcgtctggctggctggcataaatatctcactcgcaatcaaattcagccgatagcggaacgggaaggcgactggagtgccatgtccggttttcaacaaaccatgcaaatgctgaatgagggcatcgttcccactgcgatgctggttgccaacgatcagatggcgctgggcgcaatgcgcgccattaccgagtccgggctgcgcgttggtgcggacatctcggtagtgggatacgacgataccgaagacagctcatgttatatcccgccgttaaccaccatcaaacaggattttcgcctgctggggcaaaccagcgtggaccgcttgctgcaactctctcagggccaggcggtgaagggcaatcagctgttgcccgtctcactggtgaaaagaaaaaccaccctggcgcccaatacgcaaaccgcctctccccgcgcgttggccgattcattaatgcagctggcacgacaggtttcccgactggaaagcgggcagtgagcgcaacgcaattaatgtaagttagctcactcattaggcaccgggatctcgaccgatgcccttgagagccttcaacccagtcagctccttccggtgggcgcggggcatgactatcgtcgccgcacttatgactgtcttctttatcatgcaactcgtaggacaggtgccggcagcgctctgggtcattttcggcgaggaccgctttcgctggagcgcgacgatgatcggcctgtcgcttgcggtattcggaatcttgcacgccctcgctcaagccttcgtcactggtcccgccaccaaacgtttcggcgagaagcaggccattatcgccggcatggcggccccacgggtgcgcatgatcgtgctcctgtcgttgaggacccggctaggctggcggggttgccttactggttagcagaatgaatcaccgatacgcgagcgaacgtgaagcgactgctgctgcaaaacgtctgcgacctgagcaacaacatgaatggtcttcggtttccgtgtttcgtaaagtctggaaacgcggaagtcagcgccctgcaccattatgttccggatctgcatcgcaggatgctgctggctaccctgtggaacacctacatctgtattaacgaagcgctggcattgaccctgagtgatttttctctggtcccgccgcatccataccgccagttgtttaccctcacaacgttccagtaaccgggcatgttcatcatcagtaacccgtatcgtgagcatcctctctcgtttcatcggtatcattacccccatgaacagaaatcccccttacacggaggcatcagtgaccaaacaggaaaaaaccgcccttaacatggcccgctttatcagaagccagacattaacgcttctggagaaactcaacgagctggacgcggatgaacaggcagacatctgtgaatcgcttcacgaccacgctgatgagctttaccgcagctgcctcgcgcgtttcggtgatgacggtgaaaacctctgacacatgcagctcccggagacggtcacagcttgtctgtaagcggatgccgggagcagacaagcccgtcagggcgcgtcagcgggtgttggcgggtgtcggggcgcagccatgacccagtcacgtagcgatagcggagtgtatactggcttaactatgcggcatcagagcagattgtactgagagtgcaccatatatgcggtgtgaaataccgcacagatgcgtaaggagaaaataccgcatcaggcgctcttccgcttcctcgctcactgactcgctgcgctcggtcgttcggctgcggcgagcggtatcagctcactcaaaggcggtaatacggttatccacagaatcaggggataacgcaggaaagaacatgtgagcaaaaggccagcaaaaggccaggaaccgtaaaaaggccgcgttgctggcgtttttccataggctccgcccccctgacgagcatcacaaaaatcgacgctcaagtcagaggtggcgaaacccgacaggactataaagataccaggcgtttccccctggaagctccctcgtgcgctctcctgttccgaccctgccgcttaccggatacctgtccgcctttctcccttcgggaagcgtggcgctttctcatagctcacgctgtaggtatctcagttcggtgtaggtcgttcgctccaagctgggctgtgtgcacgaaccccccgttcagcccgaccgctgcgccttatccggtaactatcgtcttgagtccaacccggtaagacacgacttatcgccactggcagcagccactggtaacaggattagcagagcgaggtatgtaggcggtgctacagagttcttgaagtggtggcctaactacggctacactagaaggacagtatttggtatctgcgctctgctgaagccagttaccttcggaaaaagagttggtagctcttgatccggcaaacaaaccaccgctggtagcggtggtttttttgtttgcaagcagcagattacgcgcagaaaaaaaggatctcaagaagatcctttgatcttttctacggggtctgacgctcagtggaacgaaaactcacgttaagggattttggtcatgaacaataaaactgtctgcttacataaacagtaatacaaggggtgttatgagccatattcaacgggaaacgtcttgctctaggccgcgattaaattccaacatggatgctgatttatatgggtataaatgggctcgcgataatgtcgggcaatcaggtgcgacaatctatcgattgtatgggaagcccgatgcgccagagttgtttctgaaacatggcaaaggtagcgttgccaatgatgttacagatgagatggtcagactaaactggctgacggaatttatgcctcttccgaccatcaagcattttatccgtactcctgatgatgcatggttactcaccactgcgatccccgggaaaacagcattccaggtattagaagaatatcctgattcaggtgaaaatattgttgatgcgctggcagtgttcctgcgccggttgcattcgattcctgtttgtaattgtccttttaacagcgatcgcgtatttcgtctcgctcaggcgcaatcacgaatgaataacggtttggttgatgcgagtgattttgatgacgagcgtaatggctggcctgttgaacaagtctggaaagaaatgcataaacttttgccattctcaccggattcagtcgtcactcatggtgatttctcacttgataaccttatttttgacgaggggaaattaataggttgtattgatgttggacgagtcggaatcgcagaccgataccaggatcttgccatcctatggaactgcctcggtgagttttctccttcattacagaaacggctttttcaaaaatatggtattgataatcctgatatgaataaattgcagtttcatttgatgctcgatgagtttttctaagaattaattcatgagcggatacatatttgaatgtatttagaaaaataaacaaataggggttccgcgcacatttccccgaaaagtgccacctgaaattgtaaacgttaatattttgttaaaattcgcgttaaatttttgttaaatcagctcattttttaaccaataggccgaaatcggcaaaatcccttataaatcaaaagaatagaccgagatagggttgagtgttgttccagtttggaacaagagtccactattaaagaacgtggactccaacgtcaaagggcgaaaaaccgtctatcagggcgatggcccactacgtgaaccatcaccctaatcaagttttttggggtcgaggtgccgtaaagcactaaatcggaaccctaaagggagcccccgatttagagcttgacggggaaagccggcgaacgtggcgagaaaggaagggaagaaagcgaaaggagcgggcgctagggcgctggcaagtgtagcggtcacgctgcgcgtaaccaccacacccgccgcgcttaatgcgccgctacagggcgcgtcccattcgcca
